# Reversible control of T cell exhaustion by NR4A transcription factors revealed through targeted protein degradation

**DOI:** 10.64898/2026.02.02.701413

**Authors:** Erik Ehinger, Kristy C Perez, Emi Sanchez, Bruno Villalobos Reveles, Barsha Dash, Leo Josue Arteaga-Vazquez, Eric Johnson, Anjana Rao, Patrick G Hogan

## Abstract

Tumor-infiltrating CD8^+^ T cells (TILs) show progressive loss of effector function and upregulation of inhibitory receptors. NR4A transcription factors have emerged as key regulators of this dysfunctional state. Here we developed degron-based systems enabling rapid degradation of endogenous NR4A proteins in both mouse and primary human T cells. We demonstrate that the continuous presence of each NR4A protein is required to maintain suppression of effector cytokines and expression of co-inhibitory receptors; degradation of individual NR4A proteins rapidly restored these functional features, with each NR4A protein exerting prominent effects on distinct as well as overlapping subsets of genes and surface markers associated with effector, memory and exhaustion programs. Transcriptional profiling of phenotypically defined populations revealed both shared and unique gene programs across NR4A family members. Through CRISPR-mediated endogenous gene editing in primary human CD8^+^ T cells, we show that targeted degradation of NR4A proteins with a small molecule degrader can maintain cytokine expression and suppress inhibitory receptor expression in cells subjected to chronic stimulation, providing a framework for a powerful strategy for therapeutic intervention.

**One Sentence Summary:** Targeted degradation of endogenous NR4A proteins reveals that individual family members maintain features of T cell dysfunction through overlapping as well as non-redundant mechanisms, providing a therapeutic strategy to restore anti-tumor function.

## Introduction

Our understanding of CD8^+^ T cell responses in chronic disease settings has evolved substantially in recent years, particularly regarding the phenomenon of T cell exhaustion. CD8^+^ T cells infiltrating tumors progressively lose effector function or fail to proliferate sufficiently, instead adopting an alternative transcriptional state known as exhaustion(*1–3*). CD8^+^ T cell exhaustion is defined by diminished cytokine production, reduced or insufficient proliferation, and sustained expression of multiple inhibitory receptors(*4*). Manipulation of T cell states – exhaustion, stemness and memory – has become central to therapeutic strategies in cancer; however, despite advances in checkpoint blockade and adoptive cell therapies which form the backbone of cancer immunotherapy, most patients fail to achieve durable tumor control, highlighting the need to better understand the underlying mechanisms that establish and maintain T cell exhaustion.

The NR4A family of transcription factors, comprising NR4A1, NR4A2, and NR4A3, are encoded by immediate early genes rapidly induced upon T cell receptor engagement. Transcriptional analyses consistently identified elevated expression of all three NR4A family members in exhausted T cells across diverse settings, including mouse models of chronic infection and cancer(*4–7*) and human tumor-infiltrating lymphocytes (TILs) (*6*, *8*), suggesting their involvement in a conserved program of T cell dysfunction. Our prior work demonstrated that combined deletion of all three *Nr4a* genes in CD8^+^ CAR T cells dramatically improved anti-tumor responses by increasing cytokine production, decreasing co-inhibitory receptor expression and increasing chromatin accessibility at effector-associated regions(*6*). Recent studies have shown escalating improvements in anti-tumor effects in mice(*9*) and progressive exhaustion marker expression in human T cells undergoing chronic antigen exposure(*10*) when progressing from single to double to triple *NR4A* deletions.

We observed a hierarchy of effect sizes in our adoptive transfer studies: *Nr4a3* deletion provided the greatest anti-tumor benefit, followed by *Nr4a2*, while *Nr4a1* deletion had the smallest effect on tumor growth and survival(*6*). Other studies showed that NR4A3 reduction through knockdown or gene deletion specifically skewed acutely stimulated CD8^+^ T cells toward memory phenotypes(*8*, *11*) and improved proliferation and cytotoxicity(*8*), whereas manipulation of other family members had minimal effects. In tumor-infiltrating CAR T cells, even incomplete NR4A3 knockdown led to increased CAR T cell accumulation in tumors, reduced PD-1 expression, and enhanced cytokine production(*8*). In other studies, NR4A1 expression in TILs correlated with increased expression of TCF1, TOX, PD-1, and Ly108, but decreased expression of TIM3 and CX3CR1(*12*); NR4A1 overexpression increased TCF1^+^ TILs while *Nr4a1* deletion reduced TCF1^+^ and increased TIM3^+^ cells(*12*). However, combined deletion of *Nr4a1* and *Nr4a2* in CD8^+^ OT-I TILs restored TCF1 expression and reduced TIM3 expression(*9*), suggesting complex interactions between family members. Additionally, NR4A family members show distinct transcriptional regulation. While NFAT1 can bind all three *Nr4a* genes, NFAT signaling in the absence of AP-1—a configuration that drives a transcriptional program of T cell dysfunction which resembles CD8^+^ T cell exhaustion—preferentially induced *Nr4a2* and *Nr4a3* expression (15-fold and 50-fold increases, respectively) while having a minimal effect on *Nr4a1*(*13*, *14*). Consistent with this, only *Nr4a2* and *Nr4a3* transcription was sensitive to calcineurin inhibitors (cyclosporin A and FK506) during acute TCR activation, whereas *Nr4a1* transcription was insensitive(*14*).

Here, we explored NR4A function by developing degron-based systems that enabled acute degradation of each NR4A proteins in both mouse and primary human T cells. Our goal was to systematically define the individual contributions of each family member and assess whether individual NR4A proteins acted to initiate overlapping or distinct aspects of the exhaustion program which was then maintained by other factors, or whether the effects of the NR4As on exhaustion-related gene expression in T cells required the ongoing presence of each protein. Using enforced expression in genetically defined mouse CD8^+^ T cells coupled with conditional degradation, we show that effector cytokine suppression and co-inhibitory receptor expression both require the continuous presence of NR4A proteins. By flow cytometry, individual NR4A proteins induced largely overlapping sets of effector and regulatory markers, with some quantitative differences, both in cells expressing or lacking the other NR4A proteins. Bulk RNA-sequencing (RNA-seq) of sorted PD-1^lo^TIM3^-^ and PD-1^hi^TIM3^+^ T cell subsets, showed that while some effector, memory and exhaustion-associated genes were induced equivalently by all three individual NR4As, NR4A2 and NR4A3 induced largely overlapping sets of genes while others were differentially induced by NR4A1. Finally, we show that targeted protein degradation of NR4A proteins, endogenously engineered to be susceptible to a small molecule degrader in human cells, can maintain cytokine expression and suppress inhibitory receptor expression even in cells subjected to chronic stimulation. Our data provide insight into the transcriptional programs induced by individual NR4A family members, indicate that individual NR4A proteins maintain distinct features of T cell dysfunction through partially overlapping and partially non-redundant mechanisms, and suggest an innovative framework for therapeutic intervention through degradation of one or more NR4A proteins.

## Results

### NR4A degradation leads to recovery of cytokine expression and reduced co-inhibitory receptor expression in murine cells

In our previous study(*6*), retroviral expression of individual NR4A proteins *in vitro* resulted in increased expression of co-inhibitory receptors and decreased cytokine production upon re-stimulation. To determine whether these effects required the ongoing presence of each NR4A protein, we developed a chemical-genetic system allowing for acute depletion of individual NR4A proteins (**Fig. S1A**). Each murine NR4A protein was fused to a mutant FKBP12 protein (FKBP12^F36V^), which recruits the cereblon E3 ligase complex in the presence of the small molecule dTAG-13, enabling proximity-induced ubiquitylation of the NR4A fusion protein followed by proteasomal degradation(*15*). In both murine CD8^+^ T cells (**Fig. 1A, B**) and HEK cells (**Fig. S1B,C**), treatment with dTAG13 resulted in time and concentration-dependent degradation, with complete depletion achieved within 4 hours at 500 nM dTAG-13.

**Figure 1.**
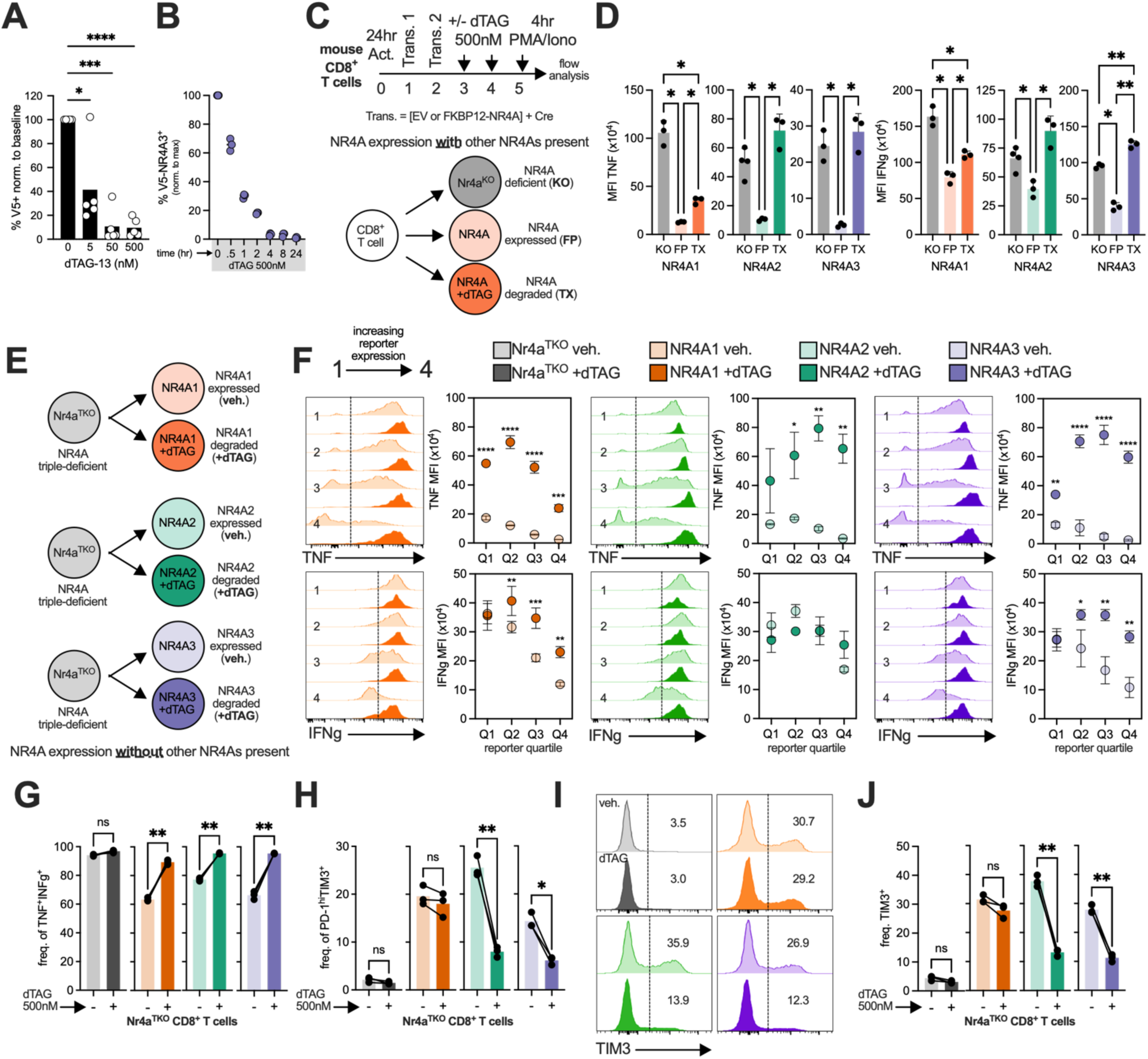
Acute depletion of individual NR4A proteins diminishes several features of T cell exhaustion. FKBP12^F36V^-V5-NR4A3 protein abundance in wildtype CD8^+^ T cells treated with (**A**) increasing concentration of dTAG-13 or (**B**) 500nM dTAG-13 for the indicated time. (**C**) Experimental scheme for reconstitution of cells lacking a single NR4A protein with the corresponding FKBP12^F36V^-NR4A fusion proteins. CD8^+^ T cells were isolated from wild-type (C57BL/6), *Nr4a1^fl^*^/fl^, *Nr4a2^fl^*^/fl^, or *Nr4a3*^-/-^ mice and retrovirally transduced with Cre and an empty vector or an expression plasmid for FKBP12^F36V^-NR4A fusion proteins; KO = Nr4a-deficient; FP = FKBP12^F36V^-NR4A; TX = FKBP12^F36V^-NR4A-expressing cells treated with dTAG-13. (**D**) MFI of single *Nr4a*-deficient CD8^+^ T cells transduced to express the indicated NR4A protein, treated (*filled symbols*) or untreated (*open symbols*) with dTAG-13, then stimulated with PMA and ionomycin for 4 hr. NR4A1, *orange*; NR4A2, *green*; NR4A3, *purple*. (**E**) Experiment graphic for reexpression of FKBP12^F36V^-NR4A fusion proteins in *Nr4a*^TKO^ cells lacking all three NR4A proteins. CD8^+^ T cells were isolated from *Nr4a1^fl^*^/fl^ *Nr4a2^fl^*^/fl^ *Nr4a3*^-/-^ (*Nr4a*^TKO^) mice and retrovirally transduced with Cre and either an empty vector or an expression plasmid for FKBP12^F36V^-NR4A fusion proteins, then treated with vehicle or dTAG-13 for 48 hours. (**F**) Reporter levels for each NR4A fusion protein expression plasmid were binned into quartiles (∼25% each) and TNF/IFNg MFI was calculated within each bin; representative histograms are shown. Reporter plasmids encoded FKBP12^F36V^-HA-NR4A1-IRES-Thy1.1 for NR4A1, FKBP12^F36V^-Flag-NR4A2-IRES-Thy1.1 for NR4A2, FKBP12^F36V^-V5-NR4A3-IRES-Thy1.1 for NR4A3. (**G**) Frequencies of TNF^+^ IFNg^+^ CD8^+^ *Nr4a*^TKO^ T cells transduced to express NR4A fusion proteins and treated or untreated with dTAG-13 for 48 hours, then stimulated with PMA and ionomycin for 4 hr. (**H**) Baseline frequencies of PD-1^hi^TIM3^+^ CD8^+^ cells transduced to express NR4A fusion proteins and untreated or treated with dTAG-13 for 48 hours. (**I**) Representative histograms of TIM3 expression. (**J**) Quantification of the TIM3^+^ T cells. Data are representative of at least two experiments. Statistical significance determined by paired t-test, one-way ANOVA with Tukey’s correction for multiple comparisons, two-way ANOVA with Šídák’s correction for multiple comparisons, or paired t-test; *p<0.05, **p<0.01, **\*\*\***p<0.001, **\*\*\*\***p<0.0001.

We focused on two key hallmarks of T cell exhaustion: diminished production of effector cytokines and other effector gene products and upregulation of co-inhibitory receptors. We first expressed individual FKBP12^F36V^-NR4A proteins in CD8^+^ T cells lacking the corresponding endogenous NR4A (e.g., NR4A3 in *Nr4a3*^-/-^ cells), then treated the cells with dTAG-13 for 48 hours (**Fig. 1C**) and evaluated baseline expression of inhibitory receptors as well as cytokine production after stimulation. For all three NR4A proteins, FKBP12^F36V^-NR4A protein (FP) expression suppressed TNF and IFNg production upon stimulation, while subsequent degradation mediated by dTAG-13 treatment (TX) restored both the frequency of cytokine-producing cells and cytokine levels (expressed as median fluorescence intensity, MFI) per cell (**Fig. 1D**). Notably, however, while all three NR4A proteins suppressed TNF (and to a lesser extent, IFNg) expression equivalently, degradation of NR4A1 was less effective at restoring cytokine expression compared to degradation of NR4A2 or NR4A3 (**Fig. 1D**). These results indicate that NR4A proteins actively repress cytokine expression even in the presence of the two other endogenous NR4As; degradation of NR4A1 has modest effects under these conditions but degradation of NR4A2 or NR4A3 efficiently restores cytokine expression.

To assess the individual contributions of each NR4A protein without compensation from other family members, we expressed the three NR4A proteins individually in CD8^+^ T cells disrupted for all three endogenous *Nr4a* genes (*Nr4a*^TKO^, **Fig. 1E**), and related cytokine expression to NR4A expression levels to levels of NR4A expression assessed as quartiles of reporter expression (**Fig. 1F**). NR4A-mediated repression of TNF occurred at all NR4A expression levels in T cells upon restimulation (**Fig. 1F**, *top panels, open symbols*), while IFNg production was almost completely repressed only at the highest expression levels of NR4A1 or NR4A3 (**Fig. 1F**, *bottom panels, open symbols*). Degradation of each NR4A protein resulted in strong recovery of TNF expression (**Fig. 1F**, *top panels, closed symbols*) and near-complete recovery of the frequency of dual TNF^+^IFNγ^+^ cells (**Fig. 1G**, **S1D**); NR4A3 degradation most effectively restored IFNg expression (**Fig. 1F**, *bottom panels, closed symbols*). Control *Nr4a*^TKO^ cells expressing empty vector showed negligible change in cytokine-producing cell frequency upon dTAG-13 treatment (**Fig. 1G**, **Fig. S1D**), though a slight increase in TNF signal intensity was observed that correlated with transduction reporter expression (**Fig. S1E, F**). This minimal effect is consistent with the reported specificity of dTAG-13, which unlike some other cereblon-binding molecules does not promote binding of cereblon to common IKZF proteins(*15*). Moreover, murine cereblon differs from human cereblon at a single residue, which abrogates IKZF recruitment(*16*), providing an additional safeguard against potential off-target effects through IKZF binding in this system.

We also examined co-inhibitory receptor expression. Expression of individual NR4A proteins resulted in formation of PD-1^hi^TIM3^+^ cells, and degradation of NR4A2/3 considerably reduced the frequency of this population (**Fig. 1H**). PD-1 expression was elevated by each NR4A; degradation of NR4A1 or NR4A2 decreased PD-1 expression, while NR4A3 degradation was not as effective (**Fig. S1G**). Thus PD-1 expression was more sensitive to regulation by NR4A1 and NR4A2, whereas TIM3 expression in PD-1^hi^TIM3^+^ cells was more influenced by NR4A2 and NR4A3.

Time course analysis revealed distinct temporal expression patterns for each receptor (**Fig. S2A-D**). TIM3 expression gradually increased over 8 days post-activation; when dTAG-13 treatment began on day 3, NR4A2/3 degradation produced effects within 24 hours, while the effects of NR4A1 degradation emerged by day 6 (**Fig. S2A**). While expression of each NR4A in *Nr4a*^TKO^ cells increased TIM3^+^ cell frequency, degradation of NR4A2 and NR4A3 more strikingly reduced both TIM3^+^ expression level and cell frequency compared to the slight decrease observed after degradation of NR4A1 (**Fig. 1I,J** and **Fig. S2A**). This effect persisted even when dTAG-13 treatment was initiated at later timepoints (day 5) (**Fig. S2E**), demonstrating that TIM3 expression requires the continuous presence of NR4A proteins and ruling against the alternative possibility that immediate TIM3 induction by NR4A proteins is later maintained through other means. PD-1 expression peaked on days 4-5 after activation in all groups and then waned; NR4A degradation accelerated this time-dependent decline (**Fig. S2B**). Other co-inhibitory receptors, such as TIGIT, were moderately induced by expression of individual NR4A proteins and their degradation led to notable decreases in the frequency and MFI of these markers (**Fig. S2C**), with NR4A2/3 again having a slightly more pronounced effect.

These findings demonstrate that NR4A-mediated cytokine repression and co-inhibitory receptor expression require the continuous presence of NR4A proteins, as degradation of individual NR4A proteins reversed these effects even in cells with high NR4A expression levels and when dTAG-13 treatment was initiated at later timepoints. Our data also suggest that the individual NR4A proteins have partly redundant and partly distinct effects on cytokine suppression and inhibitory receptor expression, although differences in initial expression levels or rates of degradation and resynthesis may contribute to the differences to some extent.

### Development of an endogenous NR4A degron system in human T cells

To study NR4A function under endogenous regulatory control, we developed a system for small molecule-mediated depletion of individual NR4A proteins that would provide us precise control of endogenous NR4A proteins in human T cells. We designed and screened panels of gRNAs targeting sites within ±10 base pairs of the start codon in the first coding exon of each *NR4A* gene (*NR4A1*, *NR4A2*, *NR4A3*) (**Fig. 2A,B**, gRNAs designated by fn1, fn2 etc). For each NR4A, we identified at least one gRNA that produced large numbers of insertions and deletions (indels) at the targeted site (e.g. NR4A1_fn1 for NR4A1, NR4A3_fn4 for NR4A3), indicating efficient targeting. We then designed homology-directed repair templates (HDRT), to be delivered via AAV transduction, encoding a different fluorescent protein reporter for each NR4A – sfGFP for NR4A1, mCherry for NR4A2, TagBFP for NR4A3 – flanked by P2A sequences and followed by V5-tagged FKBP12^F36V^ (**Fig. 2C**). T cells were electroporated with the appropriate Cas9/gRNA RNP complexes and subsequently transduced with AAV6 carrying an HDR template, then cultured for 8 days in high IL-2 (**Fig. 2D**). Successful integration produces a single transcript that, upon translation, yields an FBKP12^F36V^-NR4A fusion protein whose protein levels can be controlled by treatment with dTAG-13, and a separate fluorescent reporter protein that marks only those cells expressing the fusion protein

**Figure 2.**
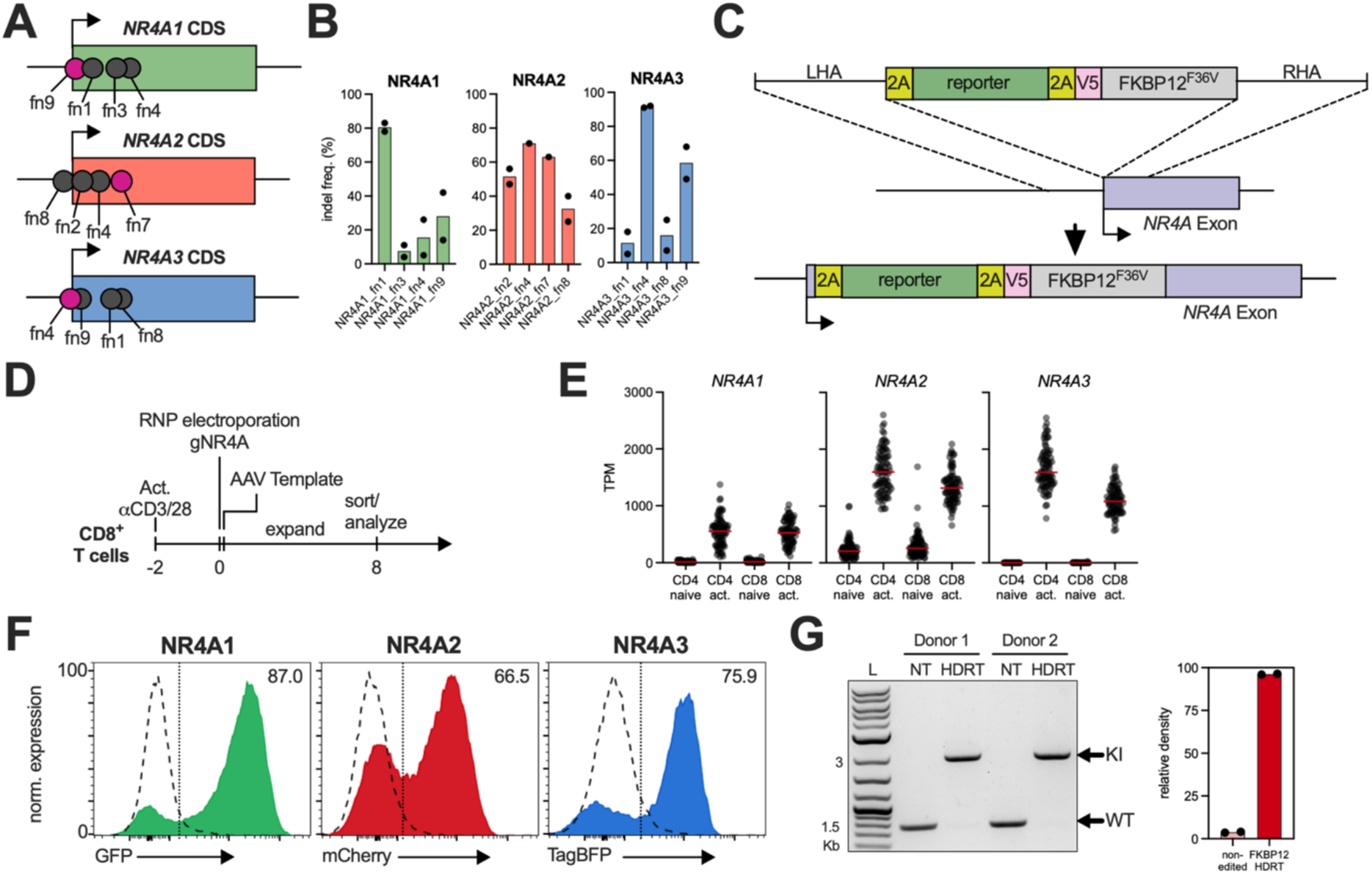
Targeted degradation of endogenous NR4A1, NR4A2, and NR4A3 differentially modulates cytokine production and exhaustion markers in human CD8^+^ T cells. **(A)** gRNAs were designed such that the cut site would be within a 10 bp proximity to the transcription start site of each NR4A locus. CD8^+^ T cells were isolated from healthy donor blood and gRNAs screened for indel frequency determined by (**B**) Inference of CRISPR Edits (ICE) analysis of each gRNA. (**C**) Schematic of HDR template delivered by AAV to achieve unique fluorescent reporter (FP) and in-frame knock-in of V5-FKBP12^F36V^ to each NR4A locus. (**D**) Experimental scheme to edit endogenous NR4As in human CD8^+^ T cells. (**E**) NR4A gene expression (TPM) in CD4 or CD8 T cells from healthy donors in the DICE Database; T cells purified based on CD3+ CD45RA+ CD127+ CCR7+ and CD4+ or CD8+, activation performed with anti-CD3/anti-CD28 beads for 4 hours. (**F**) Representative histograms of fluorescent protein expression in T cells receiving AAV-delivered HDR templates, measured by flow cytometry 4 hours after ionomycin activation. (**G**) CD8^+^ human T cells from two donors were electroporated with gRNAs targeting NR4A2 locus or a control locus (AAVS1) and subsequently delivered HDRT to insert a mCherry reporter and V5-FKBP12^F36V^. T cells were purified by mCherry^+^ reporter expression and genomic DNA isolated. PCR amplicons of NR4A2 target locus were run on a 1% agarose gel to estimate bi-allelic integration frequency based on band density (ImageJ); quantification (right). All flow cytometry data gated on live CD8^+^ events. Experiments performed in at least two independent donors.

Because endogenous NR4A proteins are low or undetectable in resting human T cells but rapidly induced upon TCR activation (from DICE database; **Fig. 2E**), fluorescent reporter expression should faithfully track endogenous NR4A expression kinetics. TCR activation (*not shown*) or ionomycin stimulation, chosen to mimic calcium-driven NFAT activation in the absence of AP-1 (**Fig. 2F**) induced robust reporter expression in HDRT-integrated cells for all three NR4A family members (**Fig. 2F**, *filled histograms*), while unstimulated cells were negative for reporter expression (**Fig. 2F**, *dashed histograms*). Fluorescent reporter expression correlated directly with integrated V5-tag levels by intracellular flow cytometry (**Fig. S3A**). Genomic PCR of reporter-positive cells revealed that the edited alleles comprised >90% of the amplification product (**Fig. 2G**, **Fig. S2B-D**), indicating bi-allelic HDRT integration in the majority of the human T cells, consistent with prior observations(*17*).

In summary, our experimental strategy yielded cells expressing all three NR4A proteins at physiological levels under endogenous control, in which the protein expression level of a single NR4A could be rapidly and reversibly regulated by targeted degradation.

### Degradation of endogenous NR4A proteins reveals partly non-redundant functions in human CD8^+^ T cells subjected to chronic stimulation

We next sought to determine how degradation of endogenous NR4A proteins affected human T cell function during chronic stimulation. Human CD8^+^ T cells were activated for 48 hours and transduced with AAV6 HDRT constructs targeting *NR4A1*, *NR4A2*, or *NR4A3*, then expanded for 14 days (**Fig. 3A,B**). On day 14, all cells were briefly stimulated with ionomycin and flow-sorted; NR4A-targeted cells were sorted based on reporter positivity, while *AAVS1*-targeted control cells, which lacked an integrated reporter, underwent the same sorting procedure. Sorted cells were immediately plated on anti-CD3/anti-CD28 antibody-coated plates with either dTAG-13 (500 nM) or vehicle control to initiate chronic stimulation. Cells were split onto fresh antibody-coated plates every 3 days as previously described(*18*). After 14 days of chronic stimulation (day 28 total), cells were either restimulated and stained for intracellular cytokines or left unstimulated and stained for extracellular markers and transcription factors (**Fig. 3A**). *AAVS1*-targeted control cells showed negligible effects of dTAG-13 treatment on all markers examined (**Fig. S4A,B**).

**Figure 3.**
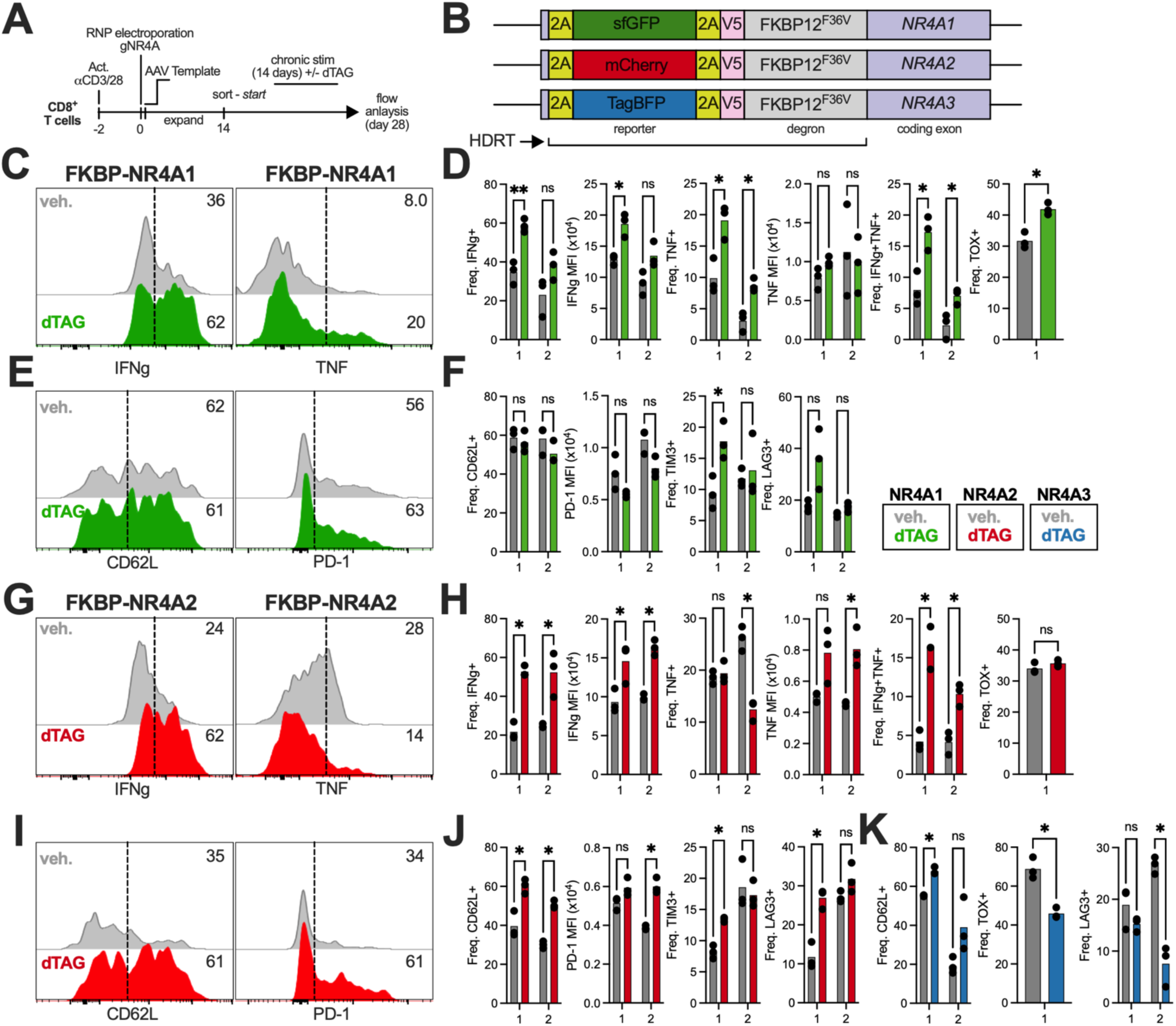
Targeted degradation of endogenous NR4A1, NR4A2, and NR4A3 differentially modulates cytokine production and exhaustion markers in human CD8^+^ T cells. **(A)** Experimental scheme for chronic stimulation analysis in CD8^+^ T cells with individually edited endogenous NR4A proteins. **(B) Illustration of HDRTs after integration into the respective *NR4A* loci. (C)** Representative histograms of IFNg (*left*) or TNF (*right*) in FKBP12^F36V^-NR4A1 edited cells treated with vehicle or dTAG-13 during chronic stimulation, then re-stimulated with ionomycin for 4 hours. **(D)** Quantification of IFNg^+^ frequency and MFI, TNF^+^ frequency and MFI, IFNg^+^TNF^+^ frequency, and TOX^+^ frequency. **(E)** Representative histograms of CD62L (*left*) or PD-1 (*right*) in FKBP12^F36V^-NR4A1 edited cells treated with vehicle or dTAG-13 during chronic stimulation. **(F)** Quantification of CD62L^+^ frequency, PD-1 MFI, TIM3^+^, and LAG3^+^ frequency. **(G)** Representative histograms of IFNg (*left*) or TNF (*right*) in FKBP12^F36V^-NR4A2 edited cells treated with vehicle or dTAG-13 during chronic stimulation, then re-stimulated with ionomycin for 4 hours. **(H)** Quantification of IFNg^+^ frequency and MFI, TNF^+^ frequency and MFI, IFNg^+^TNF^+^ frequency and TOX^+^ frequency. **(I)** Representative histograms of CD62L (*left*) or PD-1 (*right*) in FKBP12^F36V^-NR4A2 edited cells treated with vehicle or dTAG-13 during chronic stimulation. **(J)** Quantification of CD62L^+^ frequency, PD-1 MFI, TIM3^+^ frequency, and LAG3^+^ frequency. **(K)** Quantification of CD62L^+^, TOX^+^, and LAG3^+^ frequencies in FKBP12^F36V^-NR4A3 edited cells treated with vehicle or dTAG-13 during chronic stimulation. All flow cytometry data gated on live CD8^+^ events. Experiments performed in at least two independent donors with technical triplicates. Statistical comparisons by unpaired t-test with Holm-Šídák’s correction for multiple comparisons: *p<0.05, **p<0.01, ***p<0.001, ****p<0.0001.

Degradation of NR4A1 enhanced effector function across donors, increasing the frequency and MFI of IFNg^+^, TNF^+^, and IFNg^+^TNF^+^ cells compared to vehicle-treated controls (**Fig. 3C,D**). Interestingly, NR4A1 degradation increased the frequency of TOX^+^ cells while having a minimal effect on memory-associated CD62L expression (**Fig. 3D,E,F**). Among co-inhibitory receptors, NR4A1 degradation moderately reduced PD-1 levels but elevated TIM3^+^ cell frequencies (**Fig. 3E,F**), possibly through a relative increase in the expression levels of NR4A2/3 compared to NR4A1 (see below).

Degradation of NR4A2 similarly enhanced cytokine production, with elevated frequencies and MFI of IFNg^+^, TNF^+^, and double-positive cells (**Fig. 3G,H**). Unlike NR4A1, NR4A2 degradation had minimal effect on TOX expression but induced a modest increase in IL-2^+^ cells (**Fig. 3H** and **Fig. S4C,D**). NR4A2 degradation also significantly elevated memory-associated CD62L expression (**Fig. 3I,J**). However, the co-inhibitory receptor profile for diverged from NR4A1; NR4A2 degradation increased both PD-1 expression and TIM3^+^ and LAG3^+^ cell frequencies at least in one donor (**Fig. 3J**).

In contrast to NR4A1 and NR4A2, degradation of NR4A3 produced minimal changes in cytokine production, and PD-1 and TIM3 expression (**Fig. S4E**, **F**). Degradation of NR4A3 led to elevated CD62L frequency, reduced TOX^+^ cell frequency, and decreased LAG3^+^ populations (**Fig. 3K**).

Collectively, these results demonstrate that degradation of individual endogenous NR4A proteins is sufficient to restore functional features in chronically stimulated human CD8^+^ T cells, with each family member regulating a somewhat distinct subset of effector and regulatory markers depending on condition. Degradation of NR4A1 and NR4A2 enhanced cytokine production (i.e. effector function) with little contribution from NR4A3, while degradation of NR4A2 and NR4A3 led to CD62L upregulation, a feature associated with increased memory T cell formation. The effects of individual NR4A expression on TOX and co-inhibitory receptor expression were divergent: for instance, TOX and TIM3 levels were increased only after NR4A1 but not NR4A2 and NR4A3 degradation, suggesting that ongoing NR4A1 expression was selectively needed to suppress their expression.

Overall, the data suggest that at physiological levels in human T cells in which the exhaustion program has been induced by chronic stimulation, NR4A1 and NR4A2 may be the predominant NR4A proteins involved in suppression of cytokine expression, while NR4A2 and NR4A3 may contribute to suppressing CD62L^+^ memory T cell formation. Expression and suppression of co-inhibitory receptor appear to be regulated in more complex ways, potentially because our readouts involve cell surface protein expression which involves an additional layer of regulation and is often more stable than mRNA expression of the corresponding genes. The mechanisms underlying increased expression of exhaustion-associated proteins upon NR4A protein degradation are less clear, but may be indirect (*summarized in* **Table S1**).

### Individual NR4A proteins can suppress cytokine production and induce co-inhibitory receptors in mouse CD8^+^ T cells

In the experiments so far, we explored the individual functional contributions of FKBP12^F36V^-NR4A fusion proteins to cytokine and inhibitory receptor expression under conditions where expression of the NR4A fusion proteins was under dTAG-13 control. To determine how expression of individual NR4A proteins without the FKBP12^F36V^ fusion partner affected cytokine production and co-inhibitory receptor expression in mouse CD8^+^ T cells, we used strategies parallel to those in **Fig. 1**. We first compared mouse CD8^+^ T cells deleted for individual *Nr4a* genes to the same cells reconstituted with the corresponding NR4As (**Figs. 4A-F**), then repeated the experiments in CD8^+^ T cells disrupted for all three *Nr4a* genes (*Nr4a*^TKO^, **Figs. 4G-I**).

**Figure 4.**
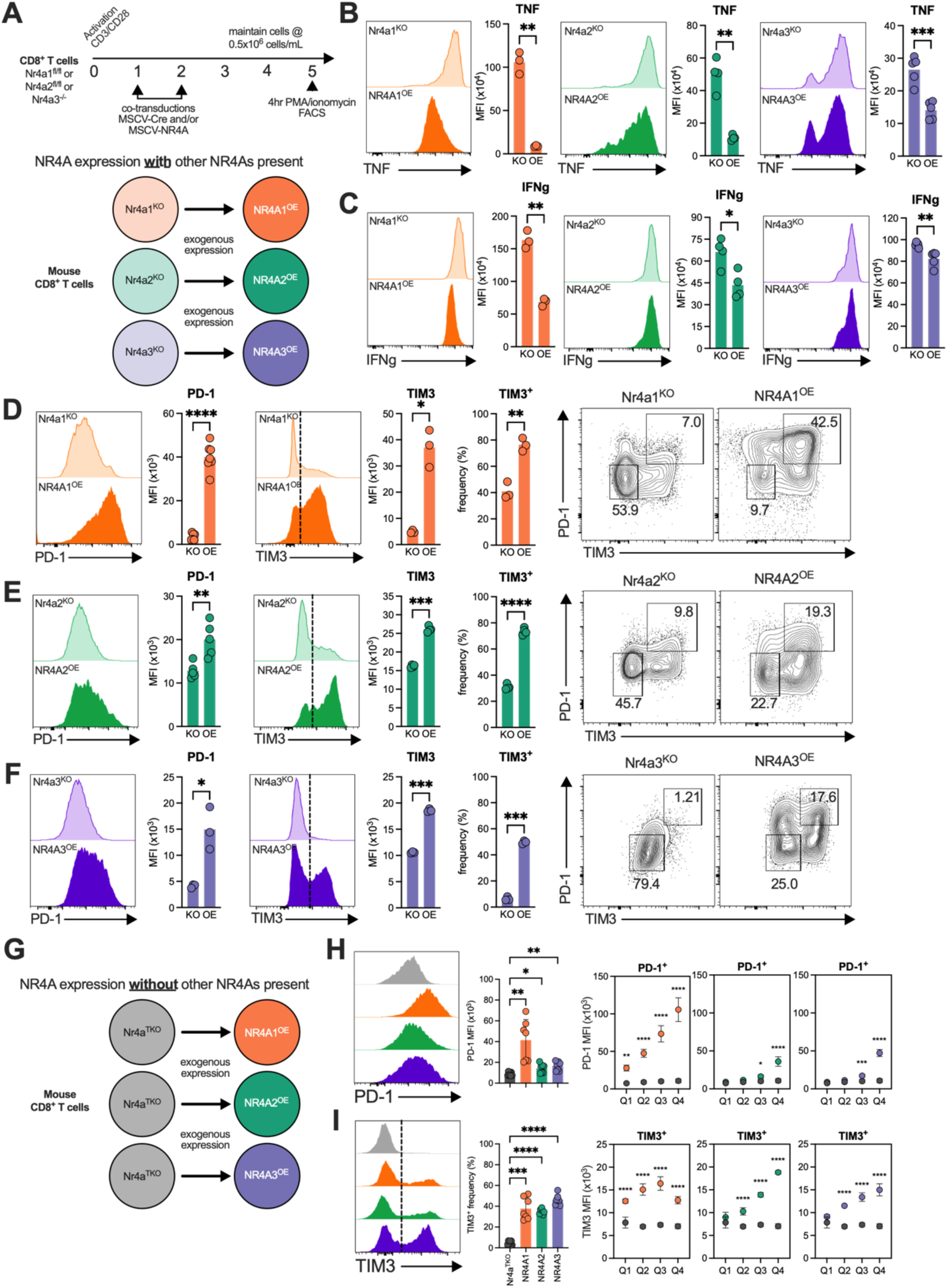
Overexpression of different NR4A proteins in single Nr4a-deficient T cells induces distinct exhaustion-associated phenotypes in vitro. **(A)** Schematic of *in vitro* CD8^+^ T cell transduction and stimulation timeline. *Nr4a1^fl/fl^, Nr4a2^fl/fl^,* or *Nr4a3^-/-^*cells were transduced to express Cre (MSCV-Cre-IRES-NGFR/GFP) and NR4A (MCSV-HA-Nr4a1- or Flag-Nr4a2- or V5-Nr4a3-IRES-NGFR/Thy1.1); *Nr4a3*^-/-^ mice did not require Cre delivery to deplete endogenous NR4A3. Representative histograms of (**B**) TNF or (**C**) IFNg expression after 4-hour PMA/ionomycin stimulation in cells gated on Cre^+^ and either empty vector or NR4A^OE^. Quantification of cytokine expression shown to the right of each histogram (n=3). Representative histograms of PD-1 (*left*) or TIM3 (*middle*) expression from unstimulated cells gated on Cre^+^ and expressing (**D**) NR4A1^OE^ (**E**) NR4A2, or (**F**) NR4A3. Quantification of surface receptor expression shown to the right of each histogram (PD-1, n=7; TIM3 n=3). *Right,* contour plots of PD-1 and TIM3 expression. (**G**) Illustration showing *in vitro* CD8^+^ T cells from *Nr4a*^TKO^ mice (*Nr4a1^fl/fl^, Nr4a2^fl/fl^,* or *Nr4a3^-/-^*) transduced to express Cre and NR4A or an empty vector. (**H**) Representative histograms of PD-1 expression in unstimulated cells (*Nr4a*^TKO^ and NR4A1/2/3^OE^) with quantification. Right, PD-1 expression across quartile bins compared to *Nr4a*^TKO^. (**I**) Representative histograms and quantification of TIM3 expression. Right, TIM3 expression across quartile bins compared to empty vector. Data is representative of at least two independent experiments. Statistical comparisons by paired T-test, one-way ANOVA with Tukey’s correction, or two-way ANOVA with Šídák’s correction: *p<0.05, **p<0.01, **\*\*\***p<0.001, **\*\*\*\***p<0.0001. PMA, Phorbol 12-myristate 13-acetate; MFI, Mean Fluorescence Intensity.

We used retroviral Cre expression to deplete individual NR4A proteins in *Nr4a1*^fl/fl^, *Nr4a2*^fl/fl^, or *Nr4a3*^-/-^ CD8^+^ T cells, and simultaneously re-expressed the depleted NR4A via retroviral expression vectors (**Fig. 4A**). Flow cytometric analysis confirmed that our expression vectors achieved NR4A protein levels comparable to the physiological range observed 2 hours post-PMA/ionomycin stimulation (**Fig. S5A**). Re-expression of each NR4A protein in *Nr4a*-depleted cells significantly suppressed both TNF and IFNg production following PMA/ionomycin stimulation (**Fig. 4B,C**). Again, more divergent results were observed for co-inhibitory receptor expression (**Figs. 4D-F**), representative histograms of PD-1 and TIM3 expression after depletion of or reconstitution with individual NR4As are shown at right. Re-expression of NR4A1 robustly induced both PD-1 and TIM3, generating populations with high expression of both receptors (**Fig. 4D**); NR4A2 re-expression increased the frequency of PD-1^+^TIM3^+^ cells but with lower MFI for both receptors, generating a substantial fraction of cells with a PD-1^lo^TIM3^+^ phenotype (**Fig. 4E**); and NR4A3 re-expression induced mixed PD-1^lo^TIM3^+^ and PD-1^+^TIM3^+^ populations characterized by broad PD-1 expression levels (**Fig. 4F**).

To assess the individual contributions of each NR4A protein without compensation from other family members, we expressed the three NR4A proteins individually in CD8^+^ T cells disrupted for all three endogenous *Nr4a* genes (*Nr4a*^TKO^) (**Fig. 4G**). Expression of individual NR4As was sufficient to decrease production of both TNF and IFNg cytokines after PMA/ionomycin stimulation (**Fig. S5B,C**). These combined effects resulted in reduced frequencies of TNF^+^IFNg^+^ double-positive cells in each case, with NR4A1 re-expression resulting in the most dramatic suppression (**Fig. S5D**). Gating on quartiles of fluorescence intensity for viral reporter expression, which correlated strongly with NR4A protein levels, we showed that high NR4A expression was associated with progressive cytokine suppression for all three NR4As (**Fig. S5E**). Notably, even cells with low NR4A levels (first quartile) showed reduced TNF expression compared to *Nr4a*^TKO^ controls, suggesting that TNF is a particularly sensitive target for NR4A-mediated repression (**Fig. S5E**, *bottom panels*). In contrast, significant IFNg suppression required higher NR4A expression levels, becoming apparent only in the third and fourth quartiles for all family members (**Fig. S5E**, *top panels*).

We next examined co-inhibitory receptor expression under conditions of individual NR4A expression. All three NR4A proteins increased PD-1 expression compared to *Nr4a*^TKO^ controls, but NR4A1 drove the most robust PD-1 induction, with expression levels strongly correlated with NR4A1 quartiles across the full range of expression (**Fig. 4H**). In contrast, NR4A2 and NR4A3 significantly increased PD-1 expression only at the highest quartiles and with substantially lower intensity (**Fig. 4H**). For TIM3, all three NR4A proteins significantly promoted increased expression (**Fig. 4I**). However, the dose-response patterns diverged: NR4A1 increased TIM3 expression progressively through the third quartile but was less effective at the highest quartile, while NR4A2 and NR4A3 exhibited stepwise increases in TIM3 with each successive quartile (**Fig. 4I**).

Collectively, these findings demonstrate that individual NR4A family members are each sufficient to suppress effector cytokine production and induce co-inhibitory receptor expression in CD8^+^ T cells, with quantitative differences in potency and dose-dependence but largely overlapping functional effects.

### NR4A family members induce distinct transcriptional programs with differential associations to progenitor and terminally exhausted T cell states

PD-1^hi^TIM3^+^ TILs are reported to have entered a late, “terminally exhausted” phase, in which the aberrant chromatin accessibility patterns associated with exhaustion are no longer reversible after adoptive transfer. To define differences in the transcriptional patterns induced by each NR4A protein in isolation from the other NR4As, we expressed each NR4A protein individually in *Nr4a*^TKO^ cells (**Fig. 5A, B**). All three NR4A proteins induced PD-1^hi^TIM3^+^ populations compared to *Nr4a*^TKO^ controls, but the individual patterns differed: NR4A1 potently induced PD-1 and TIM3 expression, enabling formation of distinct PD-1^hi^TIM3^-^ and PD-1^hi^TIM3^+^ subsets, while NR4A2 and NR4A3 more effectively increased TIM3 expression, primarily generating PD-1^hi^TIM3^+^ and PD-1^lo^TIM3^-^ populations (**Fig. 5A, B**). The pattern is similar to that shown in **Figs. 4D-F**, with the difference that other NR4As are present in the experiments of **Fig 4D-F** while only a single NR4A protein is expressed in the *Nr4a*^TKO^ background in the experiment of **Fig. 5A**.

**Figure 5.**
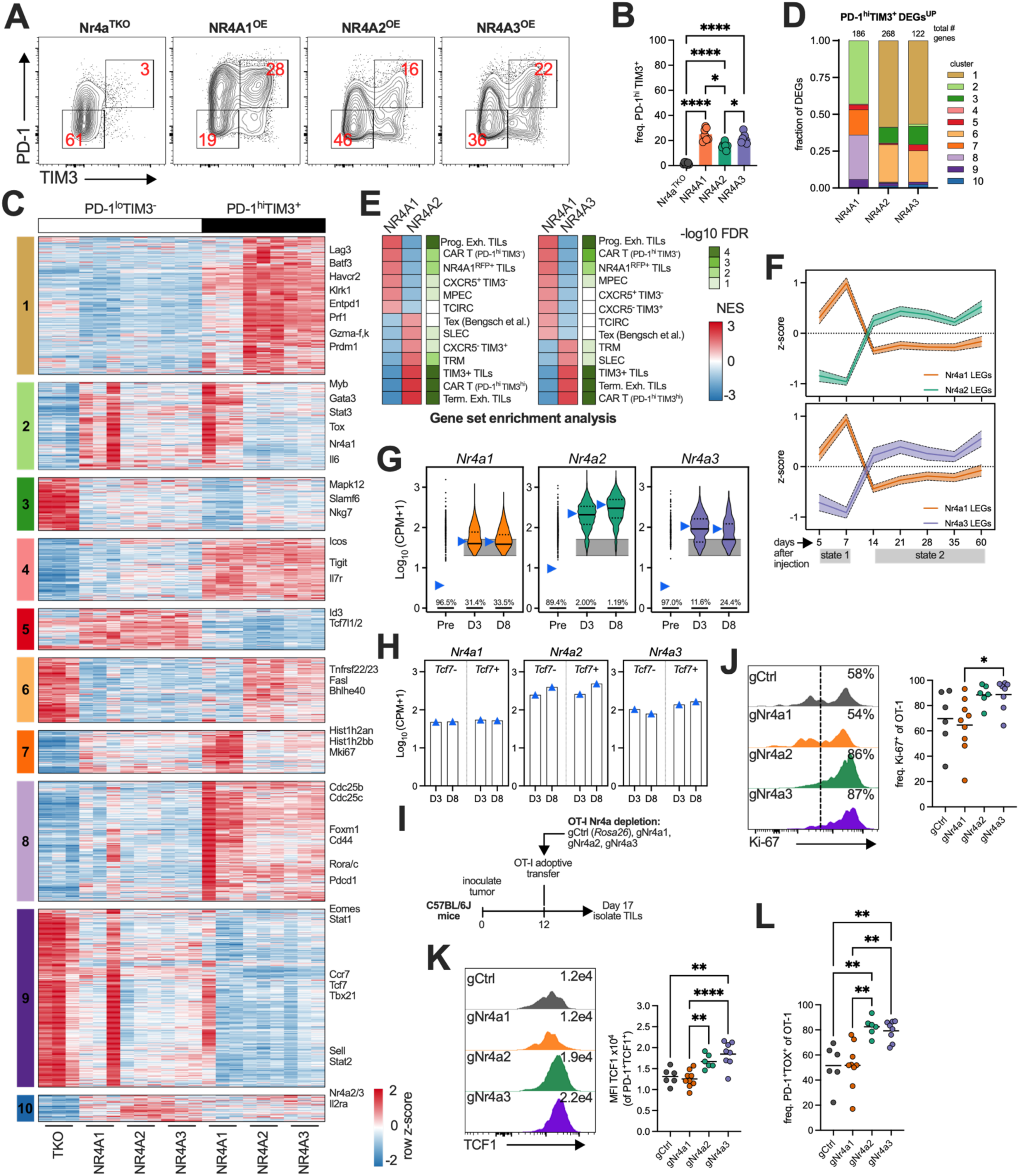
Transcriptomic analysis reveals NR4A1 promotes progenitor-like exhaustion while NR4A2 and NR4A3 induce terminal exhaustion signatures. **(A)** Representative contour plots of PD-1 and TIM3 co-expression. (**B**) Frequency of PD-1^hi^TIM3^+^ population. (**C**) K-means clustering (K=10) of z-score normalized (row) RNAseq of *Nr4a*^TKO^ CD8^+^ T cells transduced to express indicated NR4A and sorted by expression of PD-1 and TIM3; gene lists merged from differentially expressed genes as determined in all possible pairwise comparisons (adjusted p-value < 0.01 and log2 fold change ≥ 1 or ≤ -1). (**D**) Distribution of differentially expressed genes from pairwise comparisons within PD-1^hi^TIM3^+^ groups across corresponding heatmap clusters from (**C**). (**E**) GSEA displaying enrichment between NR4As sorted on PD-1^hi^TIM3^+^ (i.e., NR4A2 vs NR4A1) for signatures of indicated *in vivo* T cell populations. NES, normalized enrichment score; MPEC, memory precursor effector cell; SLEC, short-lived effector cell; TRM, tissue resident memory; TCIRC, circulating T cell. Datasets from GSE122713, GSE84105, GSE8678, GSE107395, GSE123739, and GSE266287. n=3 biological replicates. (**F**) Enrichment of leading-edge genes in murine transgenic TCR T cells that were adoptively transferred to tumor-bearing mice and harvested at the indicated timepoints from tumors; leading edge genes were normalized by z-score across days. Line is mean +/- SEM; GSE89309. State 1 dysfunction is plastic, state 2 is fixed. OT-1 CD8+ T cells, transduced with an empty vector to express GFP were adoptively transferred to CD45.1 mice 12 days after B16-OVA tumor inoculation. Pre-transfer (Pre) T cells, or TILs isolated 3 (D3), or 8 (D8) days later were subjected to single-cell RNA-sequencing (**G**) normalized (Log10(CPM+1)) counts of *Nr4a1*, *Nr4a2*, and *Nr4a3* in total cells (Pre) or cells filtered for 20k-50k total reads (D3, D8) and plotted as a violin; frequency at bottom represents the percent of cells with 0 detectable reads of each *Nr4a*; gray area marks the lowest possible expression value from a cell with 50k or 20k total read counts; blue arrows indicate pseudobulk calculation. (**H**) normalized psuedobulk (Log10(CPM+1)) counts of *Nr4a1*, *Nr4a2*, and *Nr4a3* in CD8^+^ TILs based on their expression of *Tcf7*. (**I**) Experimental design. 1.5×10^6^ OT-1 CD8^+^ T cells, modified by delivering RNPs targeting either the *Rosa26* locus (Ctrl), *Nr4a1*, *Nr4a2*, or *Nr4a3*, were adoptively transferred into CD45.1 mice 12 days after B16-OVA tumor inoculation. TILs were isolated 5 days later (day 17) and assessed by flow cytometry. (**J**) Representative histograms of Ki-67 expression in OT-1 TILs; frequency quantification (*right*). (**K**) Representative histograms of TCF1 expression in PD-1^hi^TCF1^+^ OT-1 TILs; number is median fluorescence intensity (MFI), MFI quantification (*right*). (**L**) Quantification of PD-1^hi^TOX^+^ frequency in OT-1 TILs. Data in **J-L** are representative of at least two independent experiments; gCtrl (n=6), gNr4a1 (n=9), gNr4a2 (n=6), gNr4a3 (n=8). Statistical comparisons by one-way ANOVA with Tukey’s correction for multiple comparisons (**B**, **J**-**L**): *p<0.05, **p<0.01, **\*\*\***p<0.001, **\*\*\*\***p<0.0001.

We focused on the extreme cell populations expressing the highest and lowest levels of PD-1 and TIM3 (*not shown*). Principal component analysis revealed that even the PD-1^lo^TIM3^-^ cell populations, observed primarily in cells with low NR4A expression, clustered distinctly from Nr4a^TKO^ controls (**Fig. S6A**), implying that even low-level NR4A expression substantially influences the transcriptome. PD-1^lo^TIM3^-^ transcriptomes were similar across all three NR4As, except for a moderate outlier in the NR4A1 group, while in PD-1^hi^TIM3^+^ cell populations, NR4A2 and NR4A3 transcriptomes clustered more closely together compared to NR4A1 (**Fig. S6A**).

To explore differences between the transcriptomes induced by individual NR4A proteins, we performed K-means clustering on the aggregated list of pairwise comparisons between all groups (**Fig. S6B**). We identified 10 clusters of differentially expressed genes (DEGs) across NR4A1-, NR4A2-, and NR4A3-expressing cells in both PD-1^lo^TIM3^-^and PD-1^hi^TIM3^+^ sorted populations (**Fig. 5C, Table S2,3**). Clusters 1, 6 and 10 contained genes expressed at higher levels in NR4A2/3-expressing compared to NR4A1-expressing cells (**Fig. 5D**). Cluster 1, which is predominantly upregulated in NR4A2- and NR4A3-expressing PD-1^hi^TIM3^+^ cells but considerably lower in the larger PD-1^hi^TIM3^+^ population of NR4A1-expressing cells, included the canonical exhaustion markers *Lag3, Havcr2* (TIM3), *Entpd1* (CD39), as well as the cytotoxic effector molecules *Prf1* and multiple granzymes (*Gzma-f, k*) (**Fig. 5C,D** and **Fig. S6C**). Co-expression of granzyme B and TIM3 in TILs is reported to characterize “terminally exhausted” TILs(*19*); thus the Cluster 1 signature may represent a differentially expressed gene signature of “terminal exhaustion” that is preferentially induced by NR4A2/3 compared to NR4A1. Cluster 6 contained *Bhlhe40* and *Zbtb32* genes encoding transcriptional regulators, *Ccr2* and *Ccr5* encoding chemokine receptors, *Fasl* encoding a cell-surface protein that triggers apoptosis, and *Tnfrsf* family members; these genes are also expressed in Nr4a^TKO^ controls, suggesting that they are controlled by higher levels of NR4A2 and NR4A3, but repressed by NR4A1. Cluster 10, the smallest cluster, included genes that were elevated in NR4A2/3-expressing PD-1^lo^TIM3^-^ populations, including *Il2ra*, *Nr4a2*, and *Nr4a3*.

Clusters 2 and 7 revealed genes more highly expressed in NR4A1-expressing compared to NR4A2/3-expressing cells. Cluster 2 contained NR4A1-specific genes upregulated in both PD-1^lo^TIM3^-^ and PD-1^hi^TIM3^+^ populations, including key transcription factors *Myb*, *Gata3*, *Tox*, and *Nr4a1* itself (**Fig. 5C,D**). TOX is another key exhaustion-associated transcription factor (*7*, *20–23*) that others have recently also noted is induced by NR4A1 expression(*12*). Cluster 7 contained genes induced at higher levels in NR4A1- compared to NR4A2/3-expressing PD-1^hi^TIM3^+^ cells and was enriched for proliferation markers synthesized primarily in the S and G2/M phases of the cell cycle, including several histone genes and *Mki67*.

Several clusters captured programs shared across all three NR4As. Clusters 4 and 8 captured a shared program across all three NR4As in PD-1^hi^TIM3^+^ cells: cluster 4 contained genes encoding co-stimulatory/inhibitory molecules (*Icos*, *Tigit,* and *Tnfrsf4* (OX40)), Treg-associated factor *Ikzf2* (Helios), and several histone genes and Cluster 8 included genes encoding activation markers (*Cd44*, *Cd38*, *Pdcd1*) and was enriched for cell cycle and division pathway genes (**Fig. S6D**). At least a subset of genes in Cluster 8 were significantly higher in NR4A1-expressing cells compared to NR4A2 or NR4A3 (**Fig. 5C,D** and **Fig. S6C**). Cluster 5 genes, enriched in PD-1^lo^TIM3^-^populations across all NR4As, included memory-associated transcription factor *Id3*(*24*) and TCF family member *Tcf7l2*. Cluster 3, uniquely expressed in Nr4a^TKO^ cells and containing *Nkg7* and *Slamf6*, may represent genes negatively regulated by NR4A expression. Similarly, cluster 9, containing memory-associated markers *Tcf7*, *Lef1*, *Sell*, and *Ccr7*, was highest in Nr4a^TKO^ controls with some expression in NR4A1 but minimally expressed in NR4A2/3 cells.

We performed gene set enrichment analysis (GSEA) comparing NR4A1 versus NR4A2/3 transcriptional signatures in PD-1^hi^TIM3^+^ populations against published gene sets from T cell subsets *in vivo* (**Fig. 5E**). In both comparisons (NR4A1 vs NR4A2 and NR4A1 vs NR4A3), NR4A1-induced gene signatures showed enrichment for progenitor exhausted T cells(*19*), CAR T cells (PD-1^+^TIM3^-^)(*6*), and memory precursor cells (MPECs)(*25*), whereas NR4A2-and NR4A3-induced gene signatures displayed high enrichment for terminally exhausted TILs(*19*), CAR T cells (PD-1^+^TIM3^+^)(*6*), and short-lived effector cells(*25*) (**Fig. 5E**). The enrichment of NR4A3-induced genes for short-lived effector cell signatures aligns with previous reports that NR4A3 limits the generation of memory precursor effector cells (MPECs)(*11*).

To validate these associations in an independent dataset, we examined leading-edge genes—those contributing most to the GSEA enrichment signal (**Table S4**)—in a time-resolved transcriptomics dataset of TILs from mice harboring autochthonous liver tumors driven by tamoxifen-induced activation of SV40 large T antigen (TAG)(*26*). In this model, TAG serves both to induce liver tumors and as the tumor antigen for TAG-specific transgenic T cells, which display synchronous tumor infiltration and follow a common progression from plastic dysfunction (’state 1’) to fixed “irreversible” dysfunction (’state 2’)(*26*). Analysis of the leading-edge genes revealed distinct temporal associations: NR4A1-induced leading-edge genes corresponded to the plastic, state 1, while NR4A2/3 leading-edge genes were enriched in state 2 (**Fig. 5F**).

To examine *Nr4a* expression dynamics during exhaustion progression *in vivo*, we performed single-cell RNA-seq on OT-1 CD8^+^ T cells before and after adoptive transfer into B16-OVA tumor-bearing mice. *Nr4a1*, *Nr4a2*, and *Nr4a3* were all upregulated by day 3 post adoptive transfer compared to pre-transfer cells and remained elevated in subsets of day 8 TILs (**Fig. 5G**). Between D3 and D8, *Nr4a1* remains elevated in both *Tc7^+^* and *Tcf7*^-^ populations; *Nr4a2* goes up regardless of Tcf7 expression, and *Nr4a3* selectively decreases in the *Tcf7*^-^ cells (**Fig. 5H**).

Given these distinct transcriptional associations, we next investigated how depletion of individual NR4A proteins affects the proliferation and accumulation of tumor-infiltrating lymphocytes *in vivo*. OT-1 T cells were depleted of specific NR4A proteins by electroporation with Cas9/gRNA ribonucleoprotein complexes (RNPs) targeting individual *Nr4a* genes (**Fig. S7A**), then adoptively transferred into mice bearing 12-day-established B16-OVA tumors. TILs were harvested five days post-transfer for analysis (**Fig. 5I**), a timepoint before significant differences in tumor burden were observed (**Fig. S7B**). OT-1 TILs depleted of NR4A2 or NR4A3 showed a modest increase in frequency of Ki-67^+^ cells—a marker for cycling cells(*27*)—compared to those depleted of NR4A1, though neither differed significantly from control gRNA-treated cells (**Fig. 5J**). Only NR4A2 depletion resulted in significantly greater TIL numbers in the tumor (**Fig. S7C**). Consistent with enhanced proliferative capacity, depletion of either NR4A2 or NR4A3 resulted in TILs with elevated TCF1 expression (measured as MFI in PD-1^+^TCF1^+^ cells) compared to both control and NR4A1-depleted cells (**Fig. 5K**). Additionally, NR4A2 and NR4A3 depletion increased the frequency of PD-1^+^TOX^+^ cells compared to both control and NR4A1-depleted cells (**Fig. 5L**). These results demonstrate that depletion of NR4A2 or NR4A3, but not NR4A1, enhances markers associated with proliferation and progenitor-like states in TILs.

## Discussion

The three NR4A transcription factors NR4A1, NR4A2 and NR4A3 are induced in CD8^+^ T cells by the calcium-activated transcription factor NFAT, especially in tumors and chronic viral infections where cells encounter ongoing continued antigen stimulation with little co-stimulation (*4–6*). Under these conditions, there is minimal activation of the NFAT partner AP-1 (Fos-Jun), and NFAT proteins promote an alternative transcriptional program, CD8^+^ T cell exhaustion, which limits the ability of CD8^+^ T cells to counter chronic viral infections or generate effective anti-tumor responses. Exhaustion results in upregulation of the secondary transcription factors TOX and NR4A, which directly or indirectly upregulate co-inhibitory receptors and TCR-proximal negative signaling pathways that curtail effector cytokine production and cytolytic responses(*7*, *20–23*). While it is clear that NR4A proteins are key transcription factors that drive CD8^+^ T cell exhaustion downstream of NFAT(*6*), the specific roles of each NR4A protein in promoting exhaustion are still a matter of debate.

Here we use several complementary strategies to probe the functions of each individual NR4A protein in shaping transcriptional landscapes, cytokine production patterns and cell surface phenotypes in both mouse and human CD8^+^ T cells. Through ectopic expression of each NR4A in mouse CD8^+^ T cells, either in WT cells expressing endogenous levels of the other two NR4As or in Nr4a^TKO^ cells that lack other NR4As, we found by flow cytometry that all three transcription factors suppressed TNF and IFNg production in response to stimulation and induced expression of the inhibitory receptors PD-1 and TIM3, but NR4A1 had the most striking effects (**Fig. 4**). In contrast, when we ectopically expressed FKBP12^F36V^-NR4A proteins in WT or Nr4a^TKO^ cells and compared the response to expression versus depletion of the fusion proteins by targeted degradation (**Fig. 1**), degradation of each NR4A fusion protein upregulated TNF expression and increased the frequency of TNF^+^IFNg^+^ cells (**Fig. 1G**), but there were some differences. NR4A3 degradation robustly restored IFNg and TNF production even at high levels of NR4A3 expression (**Fig. 1A-F**), and degradation of either NR4A2 or NR4A3 was more effective than NR4A1 degradation at limiting PD-1 and TIM3 expression (**Fig. 1H-J**). Using the same degradation strategy on FKBP12^F36V^-NR4A proteins expressed from the endogenous *NR4A* gene loci in human CD8^+^ T cells, NR4A1 and NR4A2 degradation led to near equivalent upregulation of cytokine production, while NR4A2 and NR4A3 degradation promoted increased expression of the memory marker CD62L. The simplest explanation for these findings is that all three NR4A proteins have similar effects on cytokine production, but the relative magnitudes of co-inhibitory receptor expression can vary depending on the experimental conditions used.

Our studies are consistent with previous comparisons of PD-1^hi^TIM3^+^ populations. NR4A1-expressing PD-1^hi^TIM3^+^ cells showed elevated *Tcf7* and *Tox* expression relative to NR4A2- or NR4A3-expressing cells, and NR4A1 bound putative regulatory regions associated with these loci(*12*). The same study showed that NR4A1 preferentially induced PD-1 over TIM3, generating distinct PD-1^hi^TIM3^-^ populations characteristic of progenitor exhausted T cells. Genes upregulated by NR4A1 were enriched in progenitor exhausted (PD-1^hi^TIM3^-^) tumor-infiltrating lymphocytes, consistent with recent findings that NR4A1 expression is highest in TIM3^-^ TILs and correlates with progenitor markers including Ly108(*12*). However, at day 3 and day 8 in tumors, all three NR4As are expressed, with the potential for complex contributions from NR4A1 as well as NR4A2/3 at early time points in TILs. Notably, NR4A2 and NR4A3 induced remarkably convergent transcriptional programs enriched for cytotoxic effector molecules (granzymes, *Prf1*) and co-inhibitory receptors associated with terminal differentiation (*Lag3*, *Havcr2*, *Entpd1*). These NR4A2/3 signatures were strongly enriched in terminally exhausted TILs, and the preferential regulation of TIM3 by NR4A2/3—evident in both overexpression and degradation experiments—further supports their role in promoting terminal exhaustion.

While cytokine expression is measured within four hours of acute antigen stimulation, the steady-state levels of cell surface protein expression at baseline after 4 days of culture are determined by many post-transcriptional and post-translational processes. We therefore turned to transcription as an acute and more reliable readout of co-inhibitory receptor expression. Because there are serious limitations to mRNA detection by single-cell RNA-seq (only a fraction of cells actually expressing a moderately abundant mRNA are actually detected as positive for that mRNA), we performed bulk RNA-seq on the cell populations expressing the highest and lowest levels of both PD-1 and TIM3. Among the *in vitro* generated CD8^+^ T cells we analyzed, the PD-1^high^TIM3^+^ cells have similar gene expression profiles to terminally exhausted CD8^+^ TILs(*6*, *12*, *19*). This population emerges at very early times after tumor infiltration and encounter with tumor antigens(*28*, *29*). Combined with K-means clustering, our strategy revealed an interesting fact that had not been as easily discerned by flow cytometry alone: that the effects of NR4A1 are often distinct from the more overlapping effects of NR4A2 and NR4A3 (**Fig. 5**). The differential effects of NR4A1 compared to NR4A2/3 on gene expression were most prominent in the PD-1^hi^TIM3^+^ population, for instance, in cluster 1 ectopic expression of NR4A2 and NR4A3 strikingly upregulated gene expression in this cluster, whereas ectopic expression of NR4A1 had almost no effect. Similarly distinct effects of NR4A1 versus NR4A2/3 were observed in cluster 2 and clusters 6-9; however, ectopic expression of any of the three NR4A proteins led to similar patterns of gene expression in clusters 3, 4, and 5 (**Fig. 5**). In line with previous studies(*6*, *10*), we conclude that depletion of all three NR4A protein represents the best strategy to alleviate exhaustion, but that the differences among NR4A family members may be exploited to fine-tune outcomes *in vivo*.

Our findings in human T cells establish targeted protein degradation as a promising modality for cancer immunotherapy. Current checkpoint blockade approaches involve treatment with antibodies targeting co-inhibitory receptors (PD-1, CTLA-4, TIM3, LAG3) that are stably upregulated in exhausted CD8^+^ T cells. These therapies have shown considerable promise in cancer patients(*30*), but blocking individual inhibitory receptors, or even combinations of multiple receptors, has not sufficed to reverse exhaustion completely, and the side effects and long-term outcomes of checkpoint blockade therapies can often be severe. Anti-PD-1 treatment facilitates the emergence of a stem-like progenitor T cell population expressing TCF-1; hence the emergence of this new population enables a new anti-tumor response rather than directly reversing exhaustion in existing T cells(*31*). In contrast, selective degradation of NR4A proteins targets upstream transcriptional programs, simultaneously reducing co-inhibitory receptor expression while restoring cytokine production and proliferative capacity—all characteristics desirable in T cell therapy. Furthermore, NR4A proteins maintain suppressive functions in multiple immune cell types; they maintain Foxp3 expression in regulatory T cells, and *Nr4a* gene deletion abrogates Treg suppressive functions(*32–36*) leading to slowed tumor growth and increased CD8^+^ TIL infiltration(*37*). In macrophages, NR4A2 directly induces anti-inflammatory M2-related genes including *Arg1* (encoding Arginase 1)(*38*), while *Nr4a1* deletion skews polarization toward proinflammatory M1-type macrophages that express higher TNF and iNOS(*39*) upon stimulation—a phenotype associated with more favorable patient outcomes(*40*). In B cells, NR4A1 depletion leads to massive proliferation among tumor-infiltrating B cells, conferring successful tumor clearance(*41*). Thus, systemic degradation of NR4A proteins, preferably by using small molecules as degraders, could both reverse T cell exhaustion and orchestrate broader anti-tumor immunity by simultaneously targeting multiple immunosuppressive cell populations.

The shared capacity for cytokine suppression among NR4A proteins likely reflects their highly conserved DNA-binding domains, which share approximately 95% amino acid sequence identity and enable binding as monomers to NBRE sites(*42*) or as homodimers to NurRE elements(*43*). Indeed, cytokine repression by NR4A1 depends entirely on DNA-binding activity; deletion or mutation of DNA-binding domains of NR4A1 abolished this function, while alterations to the ligand-binding domain had no effect(*12*). NR4A binds to regulatory regions of the *Ifng* and *Tnf* loci, supporting direct transcriptional repression of these cytokine genes(*6*, *12*). In contrast, the difference between NR4A1 and NR4A2/3 likely reflect differences in their transactivation and ligand-binding domains, which show greater sequence diversity (only ∼27-58% identity among NR4A family members)(*44*), potentially enabling differential cofactor recruitment and transcriptional output.

While depleting multiple NR4As by targeted degradation would confer more potent anti-tumor responses, it may be more practical to deplete a single NR4A protein, requiring further study of the effects of depleting a single NR4A protein in subsequent preclinical studies. There may be particular reasons to consider NR4A3 (*6*, *8*, *10*, *11*, *45*). In each of these papers, depletion (shRNA, gRNA/RNP) or gene deletion of *Nr4a3*, confers improved anti-tumor responses to varying degrees. Our own studies agree with this conclusion, except that in our models, depletion/deletion of NR4A2 and NR4A3 confer similar effects on T cell function.

In summary, our studies provide rigorous molecular evidence for the idea that targeting transcription factors that orchestrate exhaustion represents a promising frontier in cancer immunotherapy, allowing the field to move upstream of surface receptors to reprogram T cell states by manipulating their transcriptional foundations. Our findings demonstrate that manipulating NR4A protein function through targeted degradation is capable of shifting exhausted cells toward more functional, stem-like states while maintaining effector capacity. The convergence of evidence across multiple experimental systems, combined with established roles for NR4A proteins across diverse immune populations(*37*, *39*, *41*, *46*), positions NR4A-targeted degradation as a compelling strategy for enhancing anti-tumor immunity. Future integration of these tools with *in vivo* tumor models and time-resolved transcriptional profiling will refine mechanistic understanding and accelerate translation toward clinical applications that restore durable T cell function in cancer.

## Methods

### Mice

C57BL/6J, OT-1, and CD45.1 mice were purchased from Jackson Laboratories and bred in-house. Nr4a-modified mouse strains (*Nr4a1^fl/fl^*, *Nr4a2 ^fl/fl^*, and *Nr4a3*^-/-^) were obtained as previously described(*6*). These mice were crossed to create triple-deficient mice (with the delivery of Cre) as used previously(*6*). Mice between the ages of 8-12 weeks were used for experiments, and were matched for both age and sex - both male and female mice were used. All mice were bred and maintained in the animal facility at the La Jolla Institute for Immunology. All experiments and procedures were approved by the LJI Animal Care Committee.

### Construction of retroviral and AAV vectors

Cre (MSCV-Cre-IRES-NGFR), and Nr4a1, Nr4a2, Nr4a3 (MCSV-HA-Nr4a1-IRES-NGFR, MCSV-HA-Nr4a2-IRES-NGFR, MCSV-HA-Nr4a3-IRES-NGFR) were constructed as previously described(*6*). Briefly, a DNA fragment encoding Cre was PCR-amplified and cloned into MSCV-IRES-NGFR (Addgene plasmid 27489). Individually, DNA fragments encoding Nr4a1 (a gift from C.-W. J. Lio, La Jolla Institute for Immunology), Nr4a2 (Addgene Plasmid 3500), and Nr4a3 (DNASU plasmid MmCD00080978) were PCR-amplified with 5′ HA-tag and cloned into MSCV-IRES-Thy1.1. For this manuscript, all Nr4a fragments were PCR-amplified from the previously described IRES-NGFR plasmids using primers that added unique molecular tags to each Nr4a (HA-Nr4a1, Flag-Nr4a2, V5-Nr4a3) and an FKBP12 mutant (FKBP12F36V) susceptible to degradation upon dTAG-13 addition. The backbone was created by restriction digesting MSCV-IRES-mCD90.1 (Thy1.1) before both fragments were ligated by Gibson assembly. Cre cDNA was similarly PCR-amplified from MSCV-Cre-IRES-NGFR and was cloned into MCSV-IRES-GFP and MCSV-CAR-IRES-Thy1.1 upstream of the hCD19 CAR construct with a flanking P2A sequence using Gibson assembly. For AAV vectors, FKBP12F36V PCR amplified fragments with overhangs were ligated to homology arms (∼600 bp) PCR-amplified from healthy donor T cell gDNA for each endogenous NR4A locus; the resulting fragment was cloned into pAAV6 (Takara).

### Transfection

Transfections were performed in 75 cm flasks and TransIT-LT1 Transfection Reagent (Mirus Bio LLC), following manufacturer’s instructions. The Platinum-E Retroviral Packaging Cell Line, Ecotropic (PlatE) cell line was purchased from Cell BioLabs, Inc. and routinely tested for Mycoplasma. Briefly, 4.25ug pCL-Eco and 14.75ug NR4A or control vector were pre-mixed in 1.9mL OPTI-MEM (Gibco) at RT. Mirus Transfection Reagent was added dropwise. Using a P1000 the solution was gently mixed and incubated at RT for 20 min. Fresh media (DMEM + 10% FBS + 1x GlutaMAX + 1x HEPES) was added to the T-75 containing the packaging cells prior to the addition of the Transfection mix.

### Retroviral Transductions

Viral supernatant was harvested 24 after transfection, media replaced, and harvested again 48 hours after transfection. Supernatant was either filtered through a 0.45 mM filter or spun at 500xg for 5 min to remove cell debris. Retro-X concentrator (Takara Bio) was used to concentrate the supernatant overnight following the manufacturer’s recommendations. On the day of transduction, concentrated supernatant was pelleted and resuspended in T cell media (DMEM with 10% FBS, 1x GlutaMAX, 1x NEAA and 1x sodium pyruvate) containing 100U ml^-1^ IL-2 to a concentration of 10X the original volume. T cells were removed from the activation plate, counted and replated at 5×10^5^ per well. Transductions were performed in 24-well plate format, using 1.25mL of fresh TCM, 500ul of Cre vector, and 250ul of NR4A-expression (or control) vector with the addition of polybrene to a final concentration of 8mg ml^-1^. Cells were mixed with a P1000 and then centrifuged at 2000xg for 2 hr at 37C in a pre-warmed centrifuge. A second transduction is performed the following day using the 48 hr supernatant.

### Flow cytometry

T cells harvested from tissue culture were washed in 1x PBS and stained for viability (1:2000) for 10 min. Cells were washed in MACS buffer and then incubated with antibodies for surface staining (1:200). After staining, cells were washed in MACS buffer at least once and then fixed with 4% paraformaldehyde in 1xPBS for 30 min. In instances of only extracellular staining, cells were washed in MACS buffer and stored at 4C until acquisition. ICS staining was performed as described below. Samples were acquired on a Cytek Aurora, configured with 5 lasers and a 96-well plate reader. Analysis was performed using FlowJo v10 (BD Biosciences). Antibodies and reagents used for flow cytometry experiments included: eBioscience™ Fixable Viability Dye eFluor™ 780 (ThermoFisher 65-0865-18), anti-human CD8a BUV395 (BD 563796), Human TruStain FcX™ Fc Receptor Blocking Solution (Biolegend 422302), anti-human TCRa/b FITC (Biolegend 306706), anti-human TCRa/b APC (Biolegend 306717), anti-human PD-1 PE/Cy7 (Biolegend 329918), anti-human TIM3 BV711 (Biolegend 345024), anti-human LAG3 APC (Biolegend 369212), anti-human CD62L BV605 (Biolgend 304806), anti-human CD39 PE (Biolegend 328208), anti-human IFNg PE/Cy7 (BD 557643), anti-human TNF BV605 (Biolegend 502936), anti-human IL-2 (BD 563946), anti-rabbit V5 (CST 13202T), Goat anti-Rabbit IgG (H+L) Highly Cross-Adsorbed Secondary Antibody, AF647 (ThermoFisher A-21245), anti-mouse PD-1 PE/Cy7 (Biolegend 329918), anti-mouse TIM3 BV421 (Biolegend 345024), anti-mouse TNF AF647 (Biolegend 506314), anti-mouse IFNg PE/Dazzle 594 (Biolegend 505845), anti-rat CD90/mouse CD90.1 (Thy-1.1) BV711 (Biolegend 202539), anti-mouse/human Ki-67 BV711 (Biolegend 151227), anti-mouse CD69 FITC (Biolegend, 104505), anti-mouse Ly108 BV510 (BD 745073), anti-mouse CD8a BUV395 (BD 565968), anti-mouse/human REAfinity^TM^ TOX APC (Miltenyi Biotec 130-118-335), anti-mouse/human TCF1/TCF7 PE (CST C63D9), anti-mouse CD45.1 BUV737 (Invitrogen 367-0453-82), anti-mouse CD45.2 BD Horizon BUV737 (BD 612779), anti-mouse Nurr1 AF647 (Santa Cruz sc-376984), anti-mouse NOR-1 PE (Santa Cruz sc-393902), anti-mouse Nur77 AF488 (eBioscence 53-5965-82), Fixation Buffer (Biolegend 420801, eBioscience™ Foxp3 / Transcription Factor Staining Buffer Set (ThermoFisher 00-5523-00).

### Mouse T cell restimulation

Five days after T cell activation, cells were counted using a BD Accuri (BD Biosciences) and replated in a round-bottom 96 well plate at 3×10^5^ cells per well. Cells were incubated for 4 hrs in T cell media containing 10nM of PMA, 500nM of ionomycin, and 1ug ml of brefeldin A at 37C. After stimulation, cells were washed in 1xPBS and stained for viability and surface markers as described above. Cells were then fixed with 4% paraformaldehyde in 1xPBS for 30 min, washed in 1x eBioscience Perm buffer, and then stained for cytokines in 1x eBioscience Perm buffer for 30 min at a final concentration of 1:200. In some instances, where a fluorescent reporter (GFP) was used, a double fixation was performed using eBioscience Foxp3 Fix/Perm buffer for 45 min after 4% PFA. Cells were then washed once with 1x eBioscience Perm Buffer and a last wash in MACS buffer (1x PBS + 2% FBS, 2mM EDTA).

### AAV Production

AAV6-ITR containing plasmids were used for transfection of HEK293T cells along with adenovirus helper and AAV Rep-Cap plasmids using polyethylenimine (PEI MAX 40K, 1mg/ml) as the transfection reagent. Nine million 293T cells were seeded in 150 mm plates and cultured for 24 hours. Cells were then transfected with 7.5 µg of cargo vector, 7.5 µg of Rep-Cap plasmid, and 22.5 µg of adenovirus helper plasmid combined with 1.485 mL of PEI for 72 hours. Transfected cells were collected using TrypLE, resuspended in AAV lysis buffer, and lysed by three rounds of freeze/thaw cycles in a dry ice/ethanol bath. Lysate was treated with benzonase endonuclease (200U/mL) for 1 hour at 37°C, followed by addition of 5M NaCl to a final concentration of 1M for 30 minutes at 37°C. Supernatant was concentrated for 24 hours using PEG-8000. AAV was purified using iodixanol gradient centrifugation and concentrated using Amicon Ultra-15 centrifugal filters (100kDa MWCO). Vector titers were determined by qPCR after DNase I and Proteinase K treatment, using ITR-targeting primers. Relative MOI was calculated by comparison to a serial dilution of vector plasmid standard.

### AAV transduction

Human T cells were activated for 48 hours with CD3/CD28 Dynabeads (Thermo Fisher), then cultured at 1×10^6^ cells/mL in human T cell medium without FBS, accounting for anticipated 30% cell loss after electroporation. After 30 minutes rest, AAV was added at MOI 2×10^5^, ensuring total AAV volume did not exceed 20% of culture volume. Media was replaced with complete human T cell medium (with supplements) the following day, maintaining cell density at 1×10^6^ cells/mL. In cases where a CAR HDRT was delivered, negative selection was performed following manufacturer recommendation; briefly CD8^+^ T cells were resuspended in 1x MojoSort Buffer (Biolegend, day 6-8 after transduction), incubated with anti-human TCRa/b biotin (Miltenyi 130-113-529) at ∼1:50 or 10ul per 10×10^6^ cells for 15 min on ice. Cells were then washed in MojoSort buffer, incubated with MojoSort Streptavidin Nanobeads (Biolegend) on ice for 15 min, washed, and placed on a magnet for 5min at RT. Unlabeled cells were pipetted to a new tube and resuspended in TCM for continued culture.

### Human T cell isolation and gene editing

Human CD8^+^ T cells were isolated from healthy male and female donors using EasySep Human CD8^+^ T Cell Isolation Kit (STEMCELL Technologies). T cells were cultured in X-VIVO 15 media (Lonza Bioscience) supplemented with 10% FBS, 50 mM β-ME, 10 mM N-Acetyl L-Cysteine, 500 U/ml IL-2, 5 ng/ml IL-7, and 5 ng/ml IL-15. Cells were seeded at 1×10^6^ cells per ml and activated for 48 hours with CD3/CD28 Dynabeads (ThermoFisher) at a 1:1 bead:cell ratio.

For gene editing, activated T cells were electroporated with individual gRNAs (IDT), using the pulse code EH115, targeting the first coding exon of *NR4A1*, *NR4A2*, or *NR4A3*; *AAVS1* or *CD8* were used as control targets. Cells were immediately recovered by adding 80 uL of T cell media (without supplements) at 37C for 30 min. Once recovered, cells were transferred to appropriate culture vessel and resuspended in complete human TCM. After 48 hours, genomic DNA was isolated using Quick-DNA™ Miniprep Plus Kit (Zymo Research). Cut sites were PCR-amplified using Q5 Hot Start High-Fidelity Master Mix 2X (NEB), gel-purified (NEB), and analyzed by Sanger sequencing. Editing efficiency was determined using Synthego ICE Analysis, with gRNA selection based on highest indel frequency across multiple donors.

### Repetitive stimulation

CD8+ T cells edited to express individual FKBP12^F36V^-tagged NR4As were stimulated with ionomycin for 4 hours, then sorted using BD FACS Aria-I or Aria-II for the top 30% of fluorescent reporter expression. Sorted CD8+ T cells were re-suspended in X-VIVO 15 media (Lonza Bioscience) supplemented with 10% FBS, 50 mM β-ME, 10 mM N-Acetyl L-Cysteine, and containing IL-2 (40 U/mL). 150,000 cells per well were stimulated with plate-bound anti-CD3 (1 mg/mL, UCHT1) and anti-CD28 antibodies (0.5 ug/mL, CD28.2) in 96-well flat-bottom plate, in technical triplicates per donor. For chronic stimulation, cells were cultured for 14 days in a humified CO2 incubator at 37°C with and cells were split 1:2 every third day and transferred into plates freshly coated with anti-CD3 and anti-CD28 antibodies. At day 14 T cells were collected, washed, and stained for flow cytometry. For cytokine staining, cells were first re-stimulated with PMA/Ionomycin for 4 hours.

### Mouse T cell isolation and culture

CD8+ T cells were isolated from lymph nodes and spleen of mice aged 8-12 weeks using the EasySep Mouse CD8+ T Cell Isolation Kit (STEMCELL Technologies) according to manufacturer’s instructions, with the modification that we used 50% of the recommended volume. Cells were activated with plate-bound anti-CD3 (1µg/ml) and anti-CD28 (1µg/ml) antibodies in T cell media supplemented with IL-2 (100 U/ml) at 37°C and maintained at a concentration of 0.5-1×10^6^ cells/mL.

### CRISPR-mediated gene disruption in mouse T cells

For generation of *Nr4a* single-depleted T cells, activated CD8^+^ T cells were electroporated with Cas9/gRNA RNP complexes targeting *Nr4a1*, *Nr4a2*, or *Nr4a3* using the Lonza Transfection System (Lonza). Briefly, RNP complexes were assembled by incubating synthetic gRNAs (IDT) with Cas9 (Berkely Macrolab) at a 2:1 molar ratio at 37C for 15 minutes. Cells were washed with PBS, resuspended in P4 Buffer (Lonza) at 2×10^6^ cells in 14 uL (for TKO) or 19uL (for sKO), and mixed with 3ul of each RNP complex; 1ul of electroporation enhancer (IDT) was added to each reaction. Electroporation was performed using pulse code CM137. Cells were immediately recovered in DMEM without supplements for 30 min and then, if necessary, transduced with retroviral expression vectors as indicated above. Knockout efficiency was verified by flow cytometry using anti-NR4A antibodies on stimulated T cells 48 hours post-electroporation.

### Cell Sorting

Cell sorting was performed on a BD FACS Aria-I, Aria-II, or Aria-Fusion by the Flow Cytometry Core Facility at the La Jolla Institute for Immunology. Briefly, cells were sorted on lymphocyte scatter, singlets by both FSC and SSC, live cells, CD8+, and PD-1/TIM3 expression. For RNA-seq experiments, 2000 T cells from PD-1hi TIM3+ (DP) and PD-1lo TIM3- (DN) were sorted into PCR tubes containing 10.5ul of CDS Sorting Solution and immediately spun down, flash frozen, and kept at -80C for temporary storage until the start of First Strand Synthesis. Duplicates of all samples were sorted as experimental backup samples.

### Bulk RNA Sequencing

CD8 T cells sorted for their expression of PD-1 and TIM3 as described above, were thawed on ice. RNA-seq libraries were prepared using SMARTseq mRNA LP (Takara, cat# 634769) following manufacturer recommendations. Slight modifications were made to optimize cDNA LD-PCR amplification cycle number and prevent over-amplification. SYBR green concentrate was added to the cDNA LD-PCR mastermix to a final concentration of 1X upon addition to cDNA First Strand Synthesis product. 5ul of the cDNA LD-PCR mix was taken from each sample and run in a separate RT-PCR amplification on a BioRad CFX96 Real-Time PCR Detection System to determine maximal cycle number. An optimal cycle number was determined for all samples for LD-PCR using 50% of the maximum SYBR signal, -2 to -3 cycles. Subsequent protocol for purification of PCR product was adjusted to reflect the removal of 5ul from the total PCR reaction volume. D5000 or D1000 Tapestation (Agilent) Qubit (ThermoFisher) measurements were used to appropriately QC each step according to manufacturer suggestions. Libraries were pooled and quantified by the Sequencing Core at the La Jolla Institute for Immunology and sequenced on an Illumina Novaseq.

### RNA-seq analysis

Quality and adaptor trimming was performed on raw RNAseq reads using TrimGalore! v0.6.10 with default parameters. Resulting paired-end reads were aligned to mouse genome mm10 using STAR v2.7.9a. Reads aligning to annotated features were counted using the HTSeq v2.0.3 program. Differentially expressed genes were identified using the DESeq2 package v1.42.0, which was also used to normalize raw counts. For MA plots, genes with less than 10 reads were pre-filtered as an initial step. A variance Stabilizing Transformation was performed and resulting values were input to the plotPCA function of the DESeq2 package v1.42.0. Venn diagrams of common up (LFC >1) or down (LFC <1) regulated significant genes (padj<0.05) in Nr4a-overexpressing T cells compared to TKO T cells, were created using Venny (v2.1), https://bioinfogp.cnb.csic.es/tools/venny/index.html. Heatmaps were made using Morpheus (https://software.broadinstitute.org/morpheus), all pairwise comparison DEGs were used as input (LFC ≥ 1 or ≤ 1, p adj. ≤ 0.01), kmeans (k=10) clustering was performed on rows. For public datasets, RNASeq data was analyzed on GEO using GEO2R or downloaded from GEO as Fastq files using the Sra-tools v3.0.10 and mapped to human genome T2T-CHM13 using STAR v2.7.9a. Reads aligning to annotated features were counted using the featureCounts program from the Subread v.2.0.6 package. Raw counts were normalized by the TPM method.

### scRNA-seq

Tumor-infiltrating lymphocytes were isolated as described above. Pre-transfer CD8+ T cells and processed tumor single-cell suspensions were incubated with fluorescently-tagged antibodies and TotalSeq™ antibody cocktails (BioLegend, B0301, B0302, B0002). Cells were sort-purified based on viability (eBioscience™ Fixable Viability Dye eFluor™ 780, ThermoFisher 65-0865-14), CD45.1 (anti-mouse CD45.1 BV711, Biolegend 110739), CD8 (anti-mouse CD8a APC, Biolegend 100712), and GFP expression using a 100 µM nozzle into tubes containing 100% FBS. Collected cells were washed in PBS with 0.04% BSA and counted. scRNA-seq libraries were generated from 10,000 cells per condition using the Chromium Single Cell 5′ Library & Gel Bead Kit v2 (10X Genomics) according to the manufacturer’s protocol. Libraries were pooled and quantified by the Sequencing Core at the La Jolla Institute for Immunology and sequenced on an Illumina NovaSeq. Alignment, filtering, barcode counting, and UMI counting were performed using Cell Ranger v2.1.0 with default settings, and Seurat was used for further filtering. Filtering criteria included removal of cells containing excess read counts of mitochondrial genes and melanocyte genes (*Mlana*, *Dct*, *Pmel*, *Mitf*, and *Tyr*), inclusion of cells expressing *Cd8a*, and a requirement of at least 10 counts of CITE-seq CD8 antibody. Raw reads for *Nr4a1*, *Nr4a2*, and *Nr4a3* were exported using R and were additionally filtered for cells with 20,000–50,000 total reads for D3 and D8 populations, while pre-transfer cells retained all cells above the original threshold. Pseudobulk values were calculated for each condition. Data were displayed as violin plots generated in Prism v10.

### GSEA analysis

GSEA was performed using the Broad Institute Software (https://www.gsea-msigdb.org/gsea/index.jsp). Enrichment scores were determined by comparing NR4A1 versus NR4A2 or NR4A1 versus NR4A3 expression signatures against multiple publicly available datasets: GSE122713, GSE84105, GSE8678, GSE107395, GSE123739, and GSE266287. Leading edge analysis was performed to identify core genes enriched in at least 3 datasets.

### Gene ontology and pathway analysis

Gene ontology analysis was performed on cluster gene lists using Metascape (https://metascape.org/gp/index.html#/main/step1). For pathway enrichment analysis (GO Biological Processes), parameters were set to a minimum overlap of 3 genes, p-value cutoff of 0.01, and minimum enrichment score of 1.5. The top enriched pathways for each cluster are displayed.

### Tumor-infiltrating lymphocyte isolation and analysis

1.5×10^6^ OT-1 CD8^+^ T cells (CD45.1 or CD45.2 background) were adoptively transferred into C57BL/6J or CD45.1 tumor-bearing mice that had been injected subcutaneously with 1×10^5^ B16-OVA tumor cells 12 days prior. Tumors were harvested 5 days post-transfer (tumor day 17). For TIL isolation, tumors were placed in 10 mL digestion medium (RPMI supplemented with 1 mg/mL Collagenase D, 0.1 mg/mL Hyaluronidase, and 30 U/mL DNase I) in gentleMACS^TM^ C tubes (Miltenyi Biotec). Tumor dissociation was performed using a 1-minute quick dissociation program on the gentleMACS^TM^ Octo Dissociator, followed by incubation for 1 hour in a shaking incubator at 37°C and 225 rpm.

Following incubation, tube contents were filtered through a 70 μm cell strainer and centrifuged to pellet cells. The pellet was resuspended in a 40% Lymphoprep/RPMI gradient and centrifuged at 2,000xg for 1 hour. The resulting supernatant was transferred to a new 50 mL tube and centrifuged at 975xg for 20 minutes. The final cell pellet was resuspended in approximately 200 μL RPMI and transferred to a 96-well plate for flow cytometry staining and analysis.

### Statistical analysis

Statistical analysis was performed using Prism 10 (Graphpad Software). Paired analyses were used where appropriate. Statistical significance was determined by two-tailed t-tests, one-way ANOVA or Mixed-effects model with Tukey’s correction for multiple comparisons, or two-way ANOVA with Bonferroni corrections for multiple comparisons. P<0.05 was considered statistically significant.

## Acknowledgements

We would like to thank C. Kim, D. Hinz, and S. Sehic Tutusic of the Flow Cytometry Core Facility at the La Jolla Institute for Immunology for cell sorting; S. Alarcon, M. Rodrigues, and R. Wu of the Sequencing Core Facility at the La Jolla Institute for Immunology for next-generation sequencing; the Department of Laboratory Animal Care (DLAC) and the animal facility for excellent support, especially Fernando Vazquez.

## Author Contributions

E.E. and A.R. conceived and designed the study; E.E., K.C.P., E.S., and B.D. designed and performed experiments, analyzed data and composed figures; E.E. prepared sequencing libraries; B.V.R. and L.J.A.-V. analyzed sequencing data. E.S. and E. J. designed and generated viral vectors and assisted in experiments. E.E., K.C.P., A.R., and P.G.H. interpreted data. E.E. and A.R. wrote the manuscript and P.G.H. provided critical assistance, with all authors contributing to the writing and providing feedback.

## Data and materials availability

RNA sequencing data is available in the Gene Expression Omnibus (GEO) Database at GSE275301. Additional information and material related to this manuscript will be available upon reasonable request.

## Supplementary Figures

**Supplemental Figure 1.**
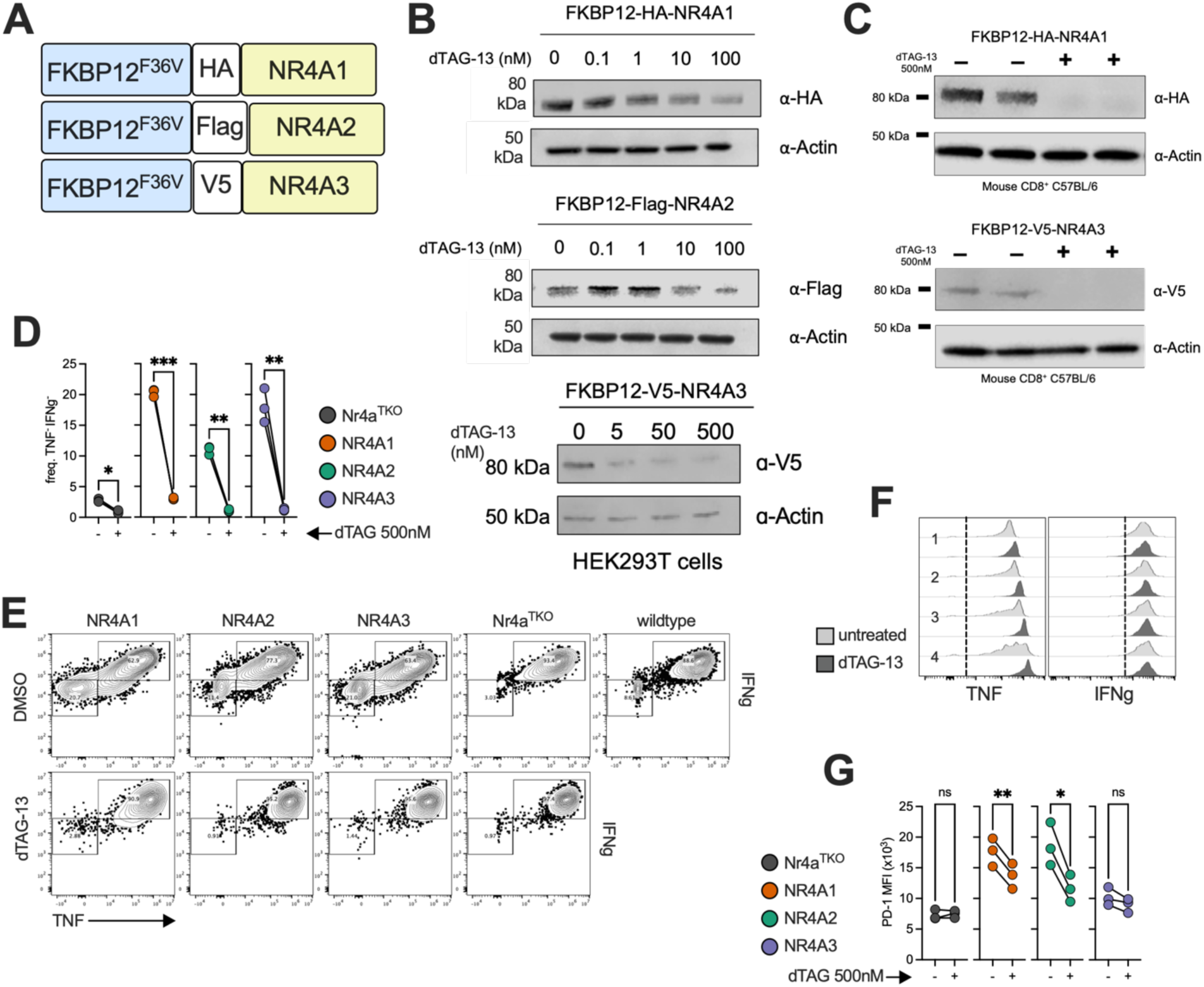
Validation of protein depletion after treatment with dTAG-13 and quantification of marker expression associated with NR4A degradation. **(A)** FKBP12^F36V^-NR4A retroviral constructs. (**B**) Western blot for associated molecular tag using HEK cell lysates after transfection with FKBP12^F36V^-NR4A constructs and treated for 4 hr with increasing doses of dTAG-13. (**C**) C57BL/6 CD8^+^ T cells were transduced with retrovirus particles encoding the FKBP12^F36V^-NR4A fusions. Three days after activation, T cells were treated with DMSO or 500nM dTAG-13 for 4 hours, lysed and assessed for protein abundance by western blot for the associated molecular tag (HA, V5); technical duplicates shown. CD8^+^ T cells were isolated from wildtype (C57BL/6) or *Nr4a1*^fl/fl^ *Nr4a2*^fl/fl^ *Nr4a3*^-/-^ mice and virally transduced with Cre and an empty vector or FKBP12^F36V^-NR4A fusion vector. Treatment was started on day 3 and analysis performed on day 5 after initial activation. (**D**) Frequencies of TNF^-^IFNg^-^ CD8^+^ *Nr4a*^TKO^ T cells transduced to express NR4A fusion proteins and treated or untreated with dTAG-13 for 48 hours, then stimulated with PMA and ionomycin for 4 hr. (**E**) representative contour plots for IFNg and TNF expression. (**F**) histograms of quartiles of reporter expression in *Nr4a*^TKO^ controls treated with DMSO or dTAG-13. (**G**) Quantification of PD-1 MFI in CD8^+^ T cells expressing FKBP12^F36V^-NR4A proteins and left untreated or treated with dTAG-13 for 48 hours. Data are representative of at least two experiments. Statistical significance determined by paired t-test; * P < 0.05, ** P < 0.01, *** P < 0.001, **** P < 0.0001.

**Supplemental Figure 2.**
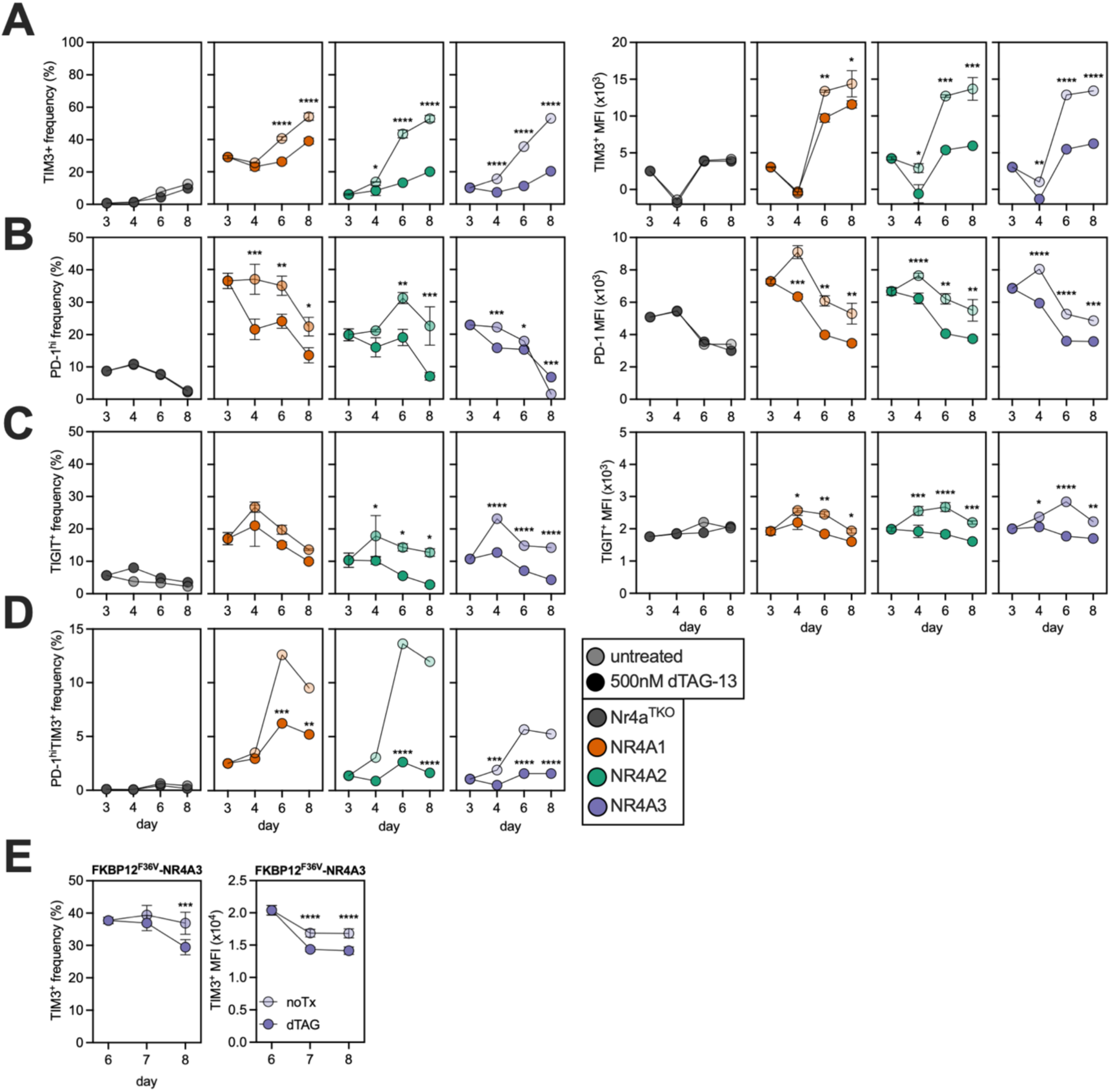
Time course analysis of marker expression after degradation of NR4As with dTAG-13 *in vitro*. Time-course analysis of indicated marker expression in CD8^+^ T cells that were isolated from wildtype (C57BL/6) or *Nr4a1*^fl/fl^ *Nr4a2*^fl/fl^ *Nr4a3*^-/-^ mice and retrovirally transduced with Cre and an empty vector or an expression plasmid for FKBP12^F36V^-NR4A fusion proteins and treated (or not) with dTAG-13 at first indicated timpoint (day); frequency (*left*), median fluorescence intensity (*right*) for (**A**) TIM3; (**B**) PD-1; (**C**) TIGIT; and (**D**) PD-1^hi^TIM3^+^. (**E**) TIM3^+^ T cell frequency (*left*) and MFI (*right*), where treatment began on day 5. Data are representative of at least two experiments. Statistical significance determined by One-way ANOVA with Tukey’s correction for multiple comparisons, or Two-way ANOVA with Sidak’s correction for multiple comparisons; * P < 0.05, ** P < 0.01, *** P < 0.001, **** P < 0.0001.

**Supplemental Figure 3.**
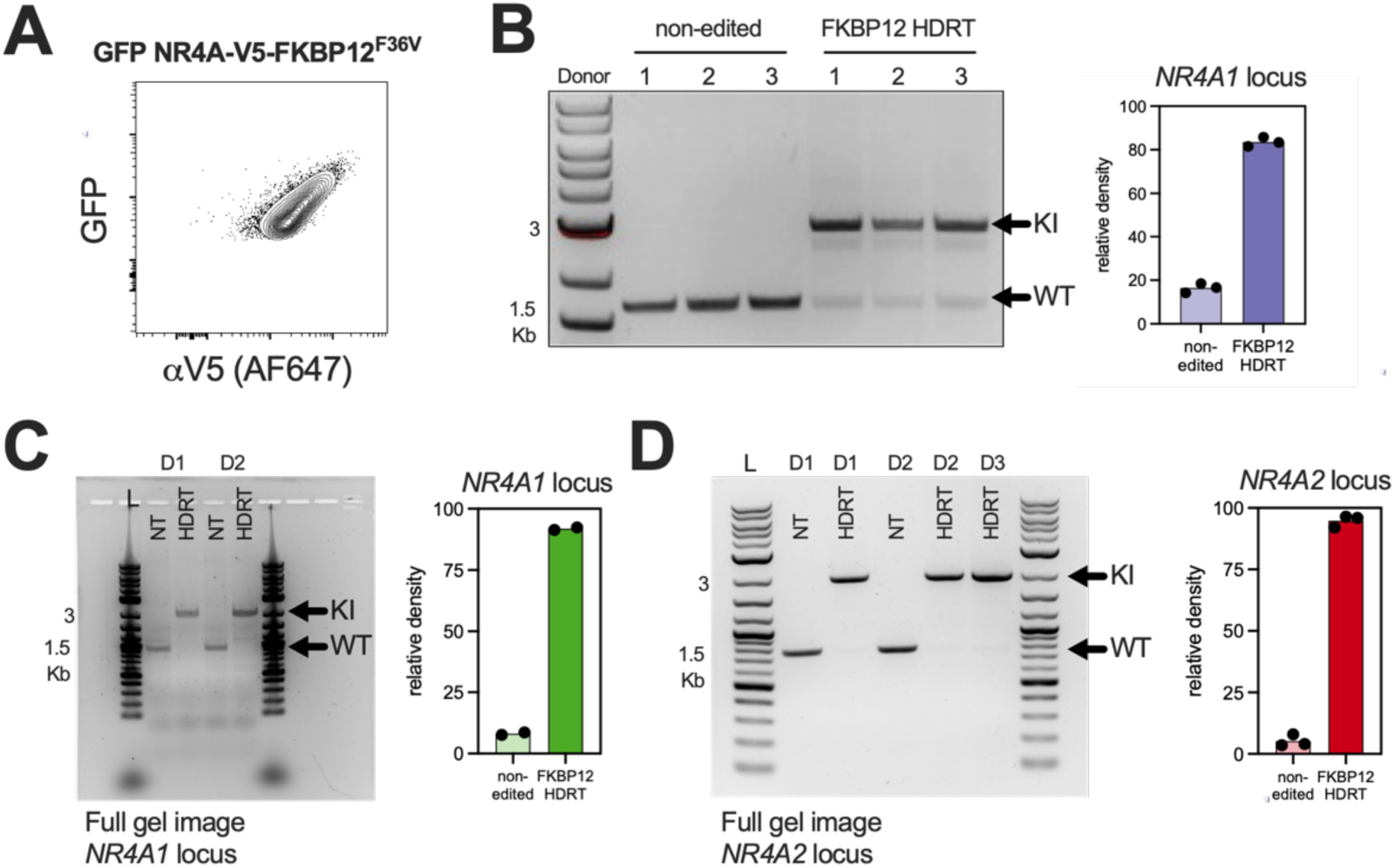
FKBP-tagged endogenous NR4A expression correlation and allelic integration. **(A)** V5-FKBP12^F36V^-NR4A1 protein abundance and GFP reporter expression by flow cytometry in human CD8^+^ T cells after 4-hour stimulation with anti-CD3/CD28 dynabeads. (**B**) CD8^+^ human T cells from three donors were electroporated with gRNAs targeting *NR4A1* locus or a control locus (*AAVS1*) and subsequently delivered HDRT to insert a GFP reporter and V5-FKBP12^F36V^. T cells were purified by GFP^+^ reporter expression and genomic DNA isolated. PCR amplicons of *NR4A1* target locus were run on a 1% agarose gel to estimate bi-allelic integration frequency based on band density (ImageJ); quantification (*right*). (**C**) Additional donors and full gel images of PCR amplicons of *NR4A1* or (**D**) *NR4A2* locus (corresponding to Fig. 6F).

**Supplemental Figure 4.**
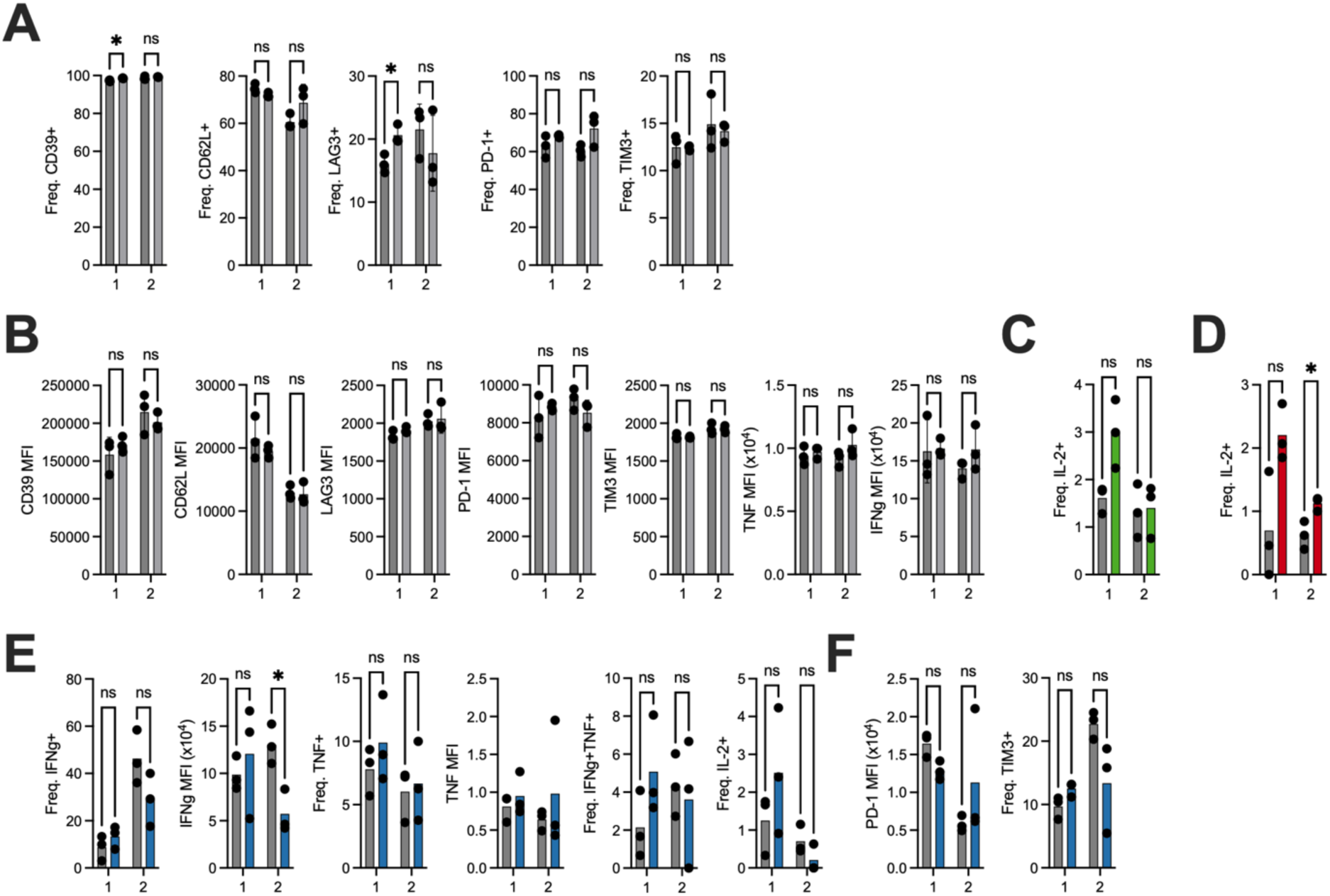
Treatment of *AAVS1*-targeted human CD8^+^ T cells or degradation of endogenous NR4A1, NR4A2, and NR4A3 with dTAG-13 during chronic stimulation. Primary human CD8^+^ T cells were electroporated with RNPs targeting the safe-harbor *AAVS1* locus, stimulated with ionomycin and sorted alongside CD8^+^ T cells expressing individually edited endogenous NR4A proteins and subjected to subsequent 14 days of chronic stimulation with anti-CD3/anti-CD28 plate-bound antibodies. (**A**) Quantification of population frequencies and (**B**) median fluorescence intensity of indicated markers between vehicle or dTAG-13 treated *AAVS1*-targeted T cells. Quantification of IL-2^+^ frequency in (**C**) FKBP12^F36V^-NR4A1 or (**D**) FKBP12^F36V^-NR4A2 edited cells treated with vehicle or dTAG-13 during chronic stimulation, then restimulated with ionomycin for 4 hours. (**E**) Quantification of IFNg^+^ frequency and MFI, TNF^+^ frequency and MFI, IFNg^+^TNF^+^ frequency in FKBP12^F36V^-NR4A3 edited cells treated with vehicle or dTAG-13 during chronic stimulation, then restimulated with ionomycin for 4 hours. Quantification of PD-1 MFI and TIM3^+^ frequency in FKBP12^F36V^-NR4A3 edited cells treated with vehicle or dTAG-13 during chronic stimulation. All flow cytometry data gated on live CD8^+^ events. Experiments performed in at least two independent donors with technical triplicates. Statistical comparisons by unpaired t-test with Holm- Šídák’s correction for multiple comparisons: *p<0.05, **p<0.01, ***p<0.001, ****p<0.0001.

**Supplemental Figure 5.**
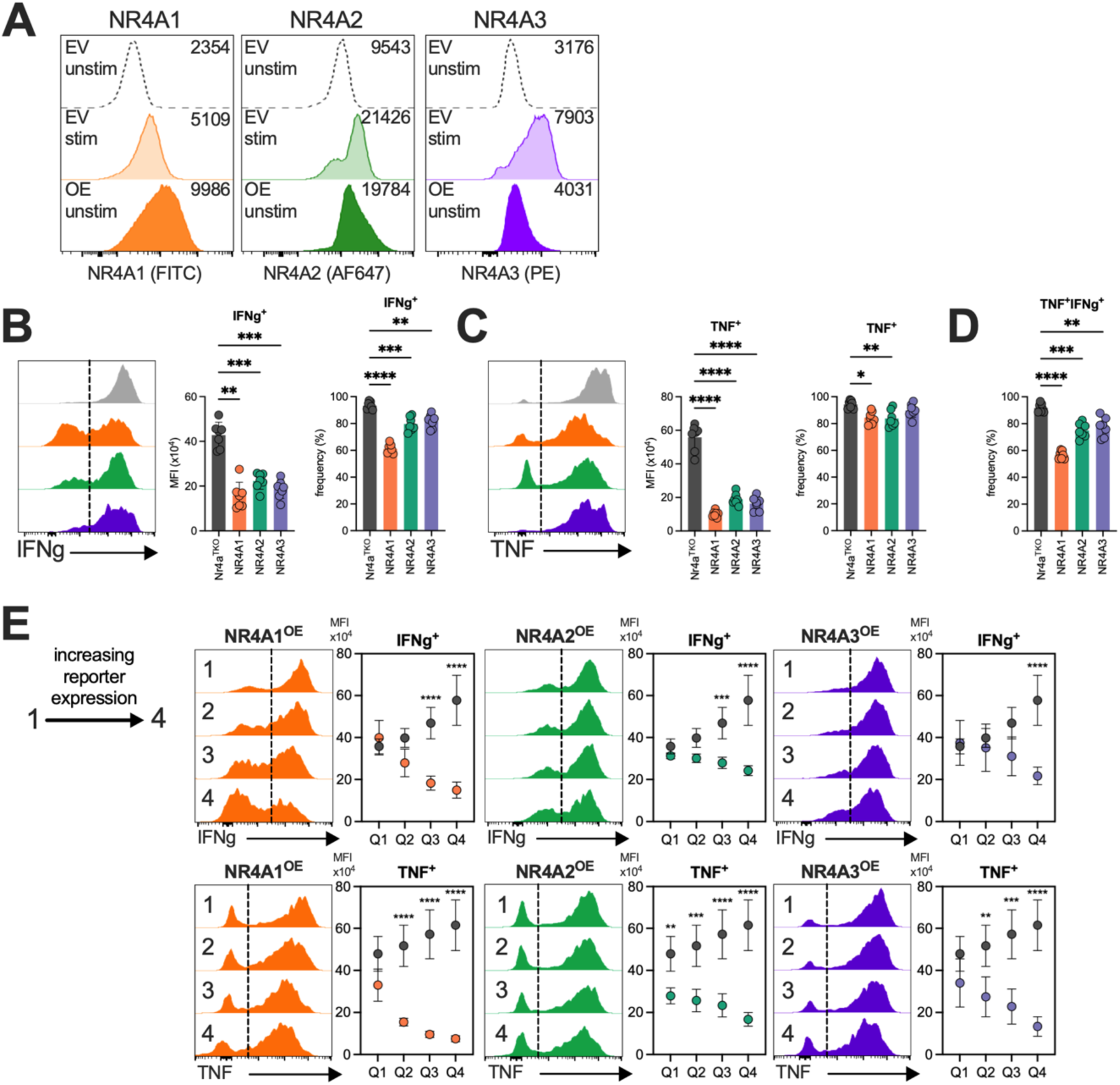
Retroviral expression of individual NR4A proteins in CD8^+^ T cells. **(A)** CD8^+^ T cells were retrovirally transduced with an empty vector (EV) or an NR4A expression vector. EV cells were stimulated for 2 hours with PMA/Ionomycin to induce expression of endogenous NR4A proteins. Numbers indicate median fluorescence intensity. CD8^+^ T cells from *Nr4a*^TKO^ mice (*Nr4a1^fl/fl^, Nr4a2^fl/fl^,* or *Nr4a3^-/-^*) were transduced to express Cre and NR4A or an empty vector. Representative histograms of (**B**) IFNg and (**C**) TNF expression from PMA/ionomycin-stimulated CD8^+^ T cells expressing empty vector (*Nr4a*^TKO^) or individual NR4A vectors (NR4A1/2/3^OE^), 5 days after initial activation. Quantification shows median fluorescence intensity (MFI) and population frequency. (**D**) Quantification of IFNg^+^TNF^+^ double-positive population frequency. For dose-response analysis (**E**), expression vector reporter levels were binned into quartiles (∼25% each, Q1-Q4), and MFI was calculated for IFNg (*top*) or TNF (*bottom*) within each bin following PMA/ionomycin stimulation. Data is representative of at least two independent experiments. Statistical comparisons by one-way ANOVA with Dunnett’s correction, or two-way ANOVA with Šídák’s correction: *p<0.05, **p<0.01, **\*\*\***p<0.001, **\*\*\*\***p<0.0001. PMA, Phorbol 12-myristate 13-acetate; MFI, Mean Fluorescence Intensity.

**Supplemental Figure 6.**
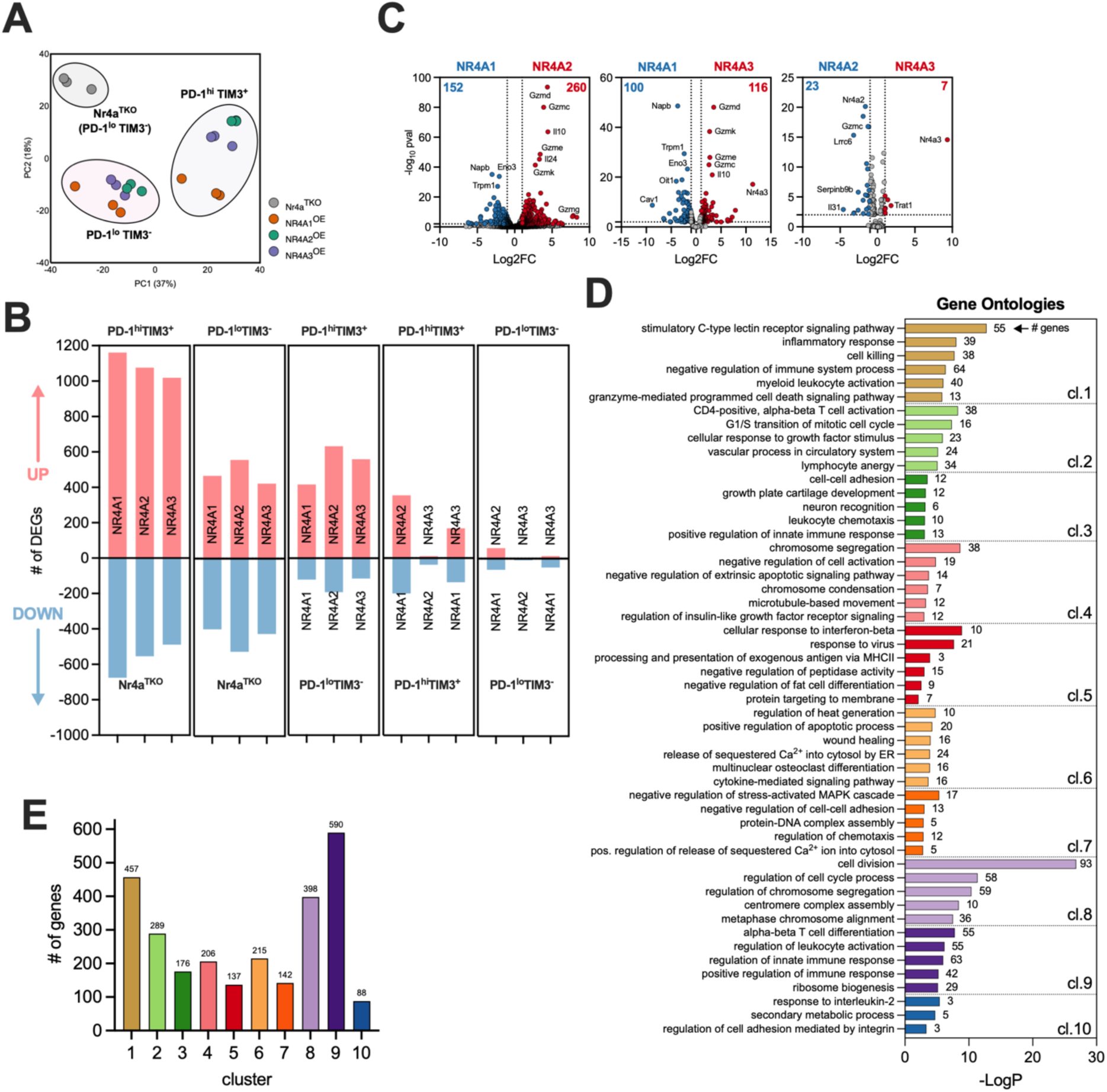
RNA sequencing of *in vitro* CD8^+^ T cells expressing individual NR4A transcription factors. CD8^+^ T cells from *Nr4a*^TKO^ mice (*Nr4a1^fl/fl^ Nr4a2^fl/fl^ Nr4a3^-/-^*) were transduced to express Cre and NR4A or an empty vector. 5 days after initial activation cells were sorted for surface marker expression of PD-1 and TIM3. (**A**) Principal component analysis (PCA) of *Nr4a* triple-deficient CD8^+^ T cells expressing empty vector (*Nr4a*^TKO^, n=3), NR4A1^OE^ (n=3), NR4A2^OE^ (n=3), or NR4A3^OE^ (n=3). (**B**) Differentially expressed genes in each indicated pairwise comparison used as input for k-means clustering. (**C**) Volcano plots of indicated pairwise comparisons between PD-1^hi^TIM3^+^ CD8^+^ T cells; significance = adjusted p-value <0.01 and log2 fold change ≥ 1 or ≤ -1. (**D**) Pathway enrichment analysis of k-means heatmap clusters in Figure 5. The top pathways are listed based on log(p-value) for each cluster, number of genes is denoted. (**E**) numbers genes within each cluster in heatmap (Fig. 5D).

**Supplemental Figure 7.**
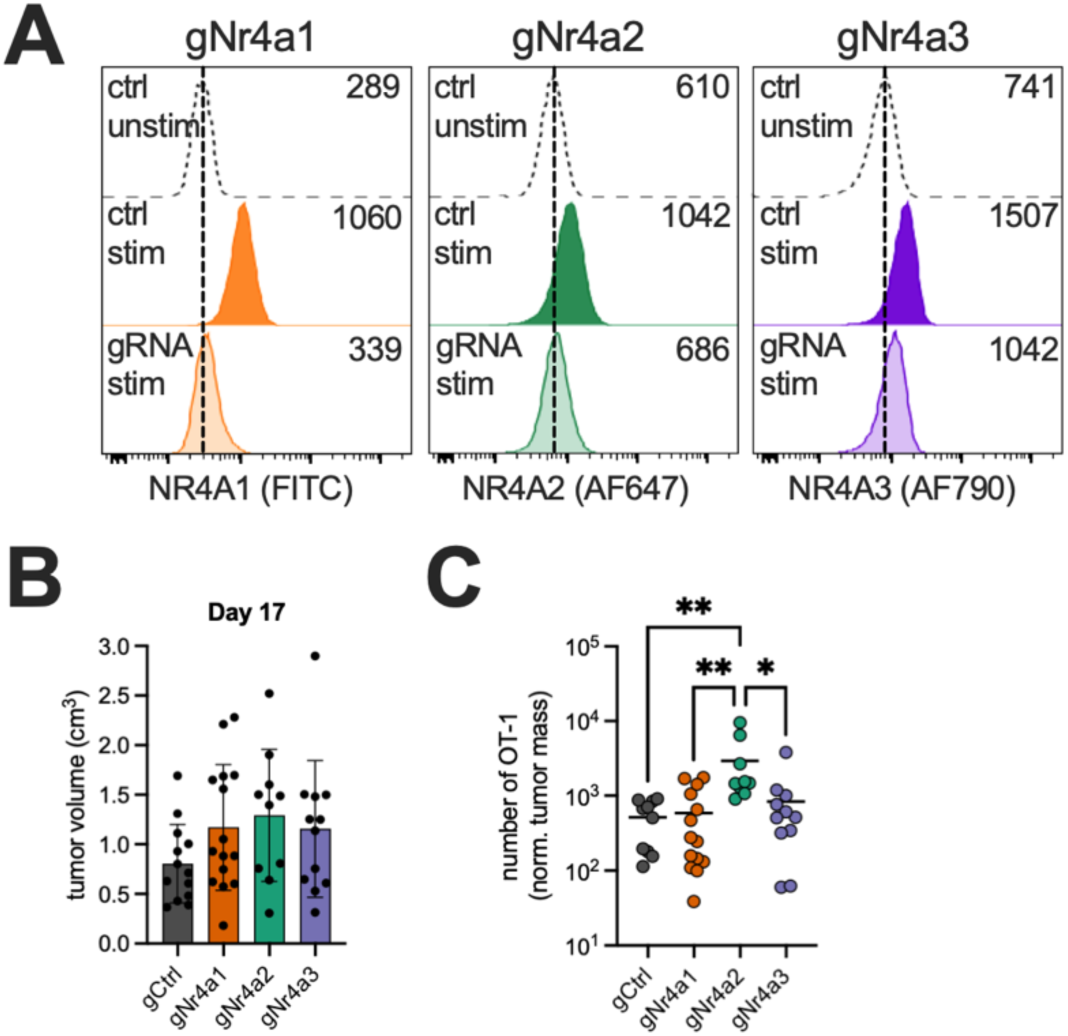
NR4A depletion effects in CD8^+^ OT-I tumor-infiltrating lymphocytes. **(A)** CD8^+^ T cells were electroporated with Cas9/gRNAs targeting individual *Nr4a* loci and evaluated for NR4A protein expression 48 hours post-electroporation following a 2-hour stimulation with PMA/Ionomycin. Numbers indicate median fluorescence intensity. (**B**) Tumor volume (cm^3^) of individual mice on tumor day 17 (5 days after OT-I adoptive transfer), measured by digital calipers; (**C**) Number of OT-1 CD8^+^ TILs normalized to tumor mass. Data in (**B-C**) are combined from two independent experiments: gCtrl (n=12), gNr4a1 (n=15), gNr4a2 (n=10), gNr4a3 (n=12); Statistical comparisons by one-way ANOVA with Tukey’s correction for multiple comparisons: *p<0.05, **p<0.01, **\*\*\***p<0.001, **\*\*\*\***p<0.0001.

